# SIRT3 preserves mitochondrial DNA integrity to restrain cGAS–STING signaling and autoimmune inflammation

**DOI:** 10.64898/2026.11.15.766791

**Authors:** Carolina Meroño, Yara Cuesta, Vanessa Marchant, Aurea Oliva, Alejandra Carrancho, Luis González-Moreno, Miguel Rodríguez-Pulido, Michael N. Sack, Margarita Sáiz, Marta Ruiz-Ortega, Javier Traba

## Abstract

Mitochondria play critical roles beyond energy production, including regulation of innate immunity through release of damage-associated molecular patterns such as mitochondrial DNA (mtDNA). Sirtuin 3 (SIRT3), a mitochondrial deacetylase, modulates mitochondrial function, reactive oxygen species (ROS) levels, and inflammation. Here, we investigate the role of SIRT3 in controlling Type I interferon (IFN) responses in macrophages.

We show that *Sirt3* depletion in mouse macrophages increases basal mtDNA release into the cytosol, activating the cGAS-STING pathway and enhancing the expression of IFN-β and downstream interferon-stimulated genes (ISGs). Accordingly, *Sirt3*-deficient cells exhibit markedly increased resistance to vesicular stomatitis virus infection. Secreted IFN-β also acts in a paracrine manner, priming neighboring cells and reinforcing antiviral defenses.

Using mtDNA-deficient (ρ^0^) cells, we demonstrate that intact mtDNA is essential for this amplified interferon response, as mtDNA depletion abolishes IFN-β induction following *Sirt3* knockdown.

Mechanistically, increased mitochondrial ROS (mtROS) in SIRT3-deficient cells promotes mtDNA leakage and cGAS-STING activation, effects that are attenuated by the mitochondrial superoxide scavenger MitoTEMPO.

*In vivo*, SIRT3 protects against chlorhexidine gluconate–induced peritoneal fibrosis by restraining macrophage infiltration, ISGs expression and fibrotic remodeling, underscoring its role in modulating STING-dependent inflammation and tissue damage. Finally, reanalysis of transcriptomic datasets from patients with lupus nephritis and rheumatoid arthritis reveals reduced SIRT3 expression, suggesting that SIRT3 downregulation is also associated with human inflammatory diseases. Collectively, our findings identify SIRT3 as an important regulator of mitochondrial integrity and oxidative stress that modulates innate immune and antiviral responses.

## INTRODUCTION

Mitochondria are essential dynamic organelles that perform diverse functions beyond adenosine triphosphate (ATP) production. They regulate heat generation, calcium (Ca²⁺) homeostasis, detoxification of reactive oxygen species (ROS), and the synthesis of heme and other key metabolites, including amino acids and nucleotides. Mitochondria also orchestrate several forms of cell death, including apoptosis, pyroptosis, ferroptosis, and necrosis. A crucial mediator in these processes is the mitochondrial permeability transition pore (mPTP), a non-specific channel in the inner mitochondrial membrane whose opening under stress conditions contributes to cell death (Traba et al., 2012).

In recent years, mitochondria have emerged as signaling organelles that significantly influence critical cellular decisions, particularly in immune system modulation (Oliva et al., 2022). Their contribution to immunity is twofold: (i) they regulate immunometabolism, determining how metabolic reprogramming affects immune cell differentiation and function (Akkaya et al., 2018; Traba et al., 2020; Traba et al., 2021; Traba & Anton, 2022); and (ii) they act as sources of damage-associated molecular patterns (DAMPs) and as platforms for innate immune signaling. During an innate immune response, they stimulate the release of pro-inflammatory cytokines and type I interferons in defense against pathogen-associated molecular patterns (PAMPs). Recent findings show that the release of mitochondrial DNA (mtDNA) into the cytosol under certain pathophysiological conditions can activate innate immune pathways, including TLR9, the NLRP3 inflammasome, and cGAS/STING signaling (West and Shadel, 2017; Sack 2018). In particular, the CGAS/STING pathway culminates with the production of type I interferons (IFNs) and interferon-stimulated genes (ISGs) which are responsible for antiviral responses (Barnett et al, 2019).

Mechanisms ensuring mitochondrial quality control, such as fission and fusion (Yasukawa et al., 2009), mitophagy (Zhou et al., 2012), and regulation by mitochondrial sirtuins (Webster et al., 2014), are essential to restrain inflammation. Sirtuins are NAD⁺-dependent deacylases that act as metabolic sensors. In mammals, the family includes seven members: SIRT1, SIRT6, and SIRT7 (nuclear); SIRT2 (cytosolic); and SIRT3, SIRT4, and SIRT5 (mitochondrial). Although most act as deacetylases, SIRT5 and SIRT4 catalyze distinct acyl-removal reactions (Du et al., 2011; Anderson et al., 2018).

Acylation in the cytosol and nucleus is governed by enzymatic processes involving lysine acetyltransferases, while mitochondrial acylation occurs non-enzymatically. This non-enzymatic acylation is influenced by higher concentrations of acyl-CoAs and the alkaline pH of the mitochondria (Ghanta et al., 2013). This distinction suggests a paradigm where mitochondrial acylation may represent protein damage—acetylated mitochondrial proteins are typically less active—, in contrast to the signaling role acylation plays in the cytosol and nucleus. SIRT3 stands out as the primary mitochondrial deacetylase and is the only sirtuin responsible for widespread mitochondrial protein deacetylation. The main SIRT3 targets are metabolic and respiratory enzymes, and this confers SIRT3 an important role in the regulation of mitochondrial function (Bause and Haigis, 2013).

We previously demonstrated that SIRT3 limits NLRP3 inflammasome activation by deacetylating and activating mitochondrial superoxide dismutase (SOD2), thereby reducing mtROS accumulation (Traba et al., 2017). SIRT3 also suppresses inflammation via additional mechanisms (Zhao et al., 2016; Liu et al., 2018; Zheng et al., 2018; Ma et al., 2018; Deng et al., 2023; Maurice et al., 2022). Moreover, NAD⁺ repletion using precursors such as nicotinamide riboside modulates inflammation through sirtuin-dependent and -independent mechanisms (Traba et al., 2015; Zhou et al., 2020; Wu et al., 2022; Doke et al., 2023; Han et al., 2023).

In this study, we explored the influence of SIRT3 on antiviral immune responses by investigating how its deficiency affects the release of mtDNA and subsequent type I interferon production. We generated macrophage models to elucidate the role of mtDNA in the antiviral innate immune response: cells with increased mtDNA leakage into the cytosol due to SIRT3 depletion and cells lacking mtDNA (ρ^0^ cells), where mtDNA leakage cannot occur. Our findings show that *Sirt3* knockdown induces a basal leak of mtDNA, which serves as a potent stimulus for type I interferon production, likely through activation of the cGAS-STING pathway. This response to mtDNA leakage confers enhanced antiviral protection, as shown by increased interferon-β release, increased expression of ISG, and heightened resistance to viral infection. Notably, we observed that cells deficient in mtDNA exhibited blunted antiviral responses when stimulated by cytosolic RNA or DNA, emphasizing that a constitutive mtDNA leak in macrophages may be critical for priming the antiviral response. These findings advance our understanding of the role of mitochondrial quality control and mtDNA in immune signaling and suggest that targeting SIRT3 may enhance antiviral responses by modulating mtDNA release and interferon signaling.

Furthermore, in a murine model of chlorhexidine-induced peritoneal injury, *Sirt3* deficiency exacerbated local inflammation and fibrotic remodeling, accompanied by enhanced macrophage infiltration and ISG expression, underscoring the physiological relevance of mitochondrial control of immune activation. In line with these findings, reanalysis of publicly available gene-expression datasets revealed reduced SIRT3 expression in glomerular samples from patients with lupus nephritis and in macrophages from patients with rheumatoid arthritis, suggesting that SIRT3 downregulation is also associated with human inflammatory diseases.

Altogether, these findings expand our understanding of how SIRT3 safeguards tissue homeostasis by restraining aberrant mtDNA-driven innate immune responses and suggest that modulating SIRT3 activity could represent a strategy to fine-tune type I interferon–dependent inflammation in disease.

## RESULTS

### Increased leak of mitochondrial DNA in SIRT3 depleted cells increases transcription of type I interferon and interferon stimulated genes

Our previous studies showed that SIRT3 depletion in primary macrophages or macrophage cell lines increases ROS levels and NLRP3 inflammasome activation (Traba et al., 2015; 2017), accompanied by elevated extrusion of mitochondrial DNA (mtDNA) into the cytosol upon stimulation with lipopolysaccharide (LPS) and ATP. Notably, SIRT3 depletion leads to a heightened basal mtDNA leakage, both in resting cells and isolated mitochondria (**Fig. 1A**).

**Figure 1.**
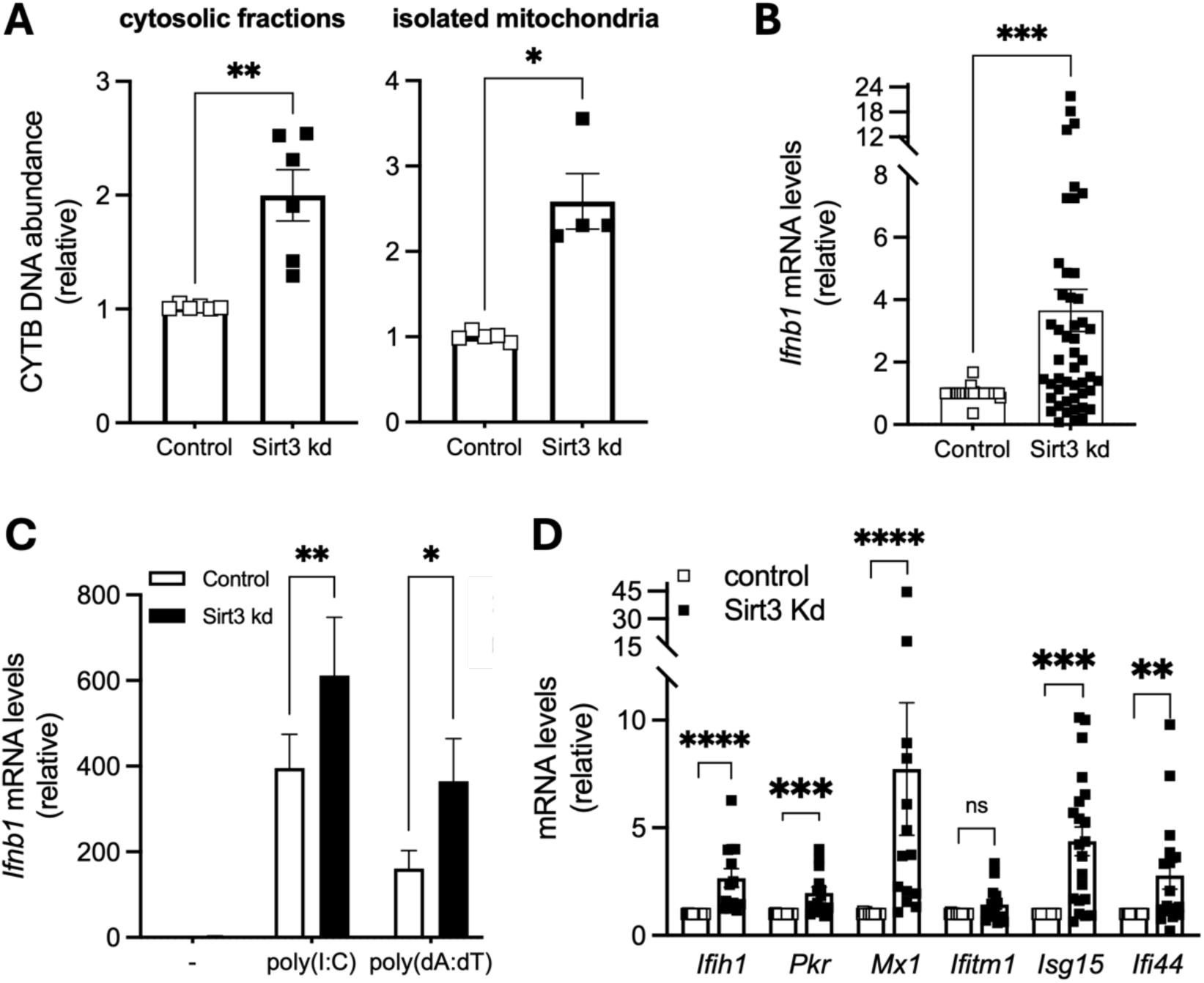
SIRT3-depleted cells exhibit increased mtDNA leak and enhanced type I interferon responses. (**A**). Release of mtDNA into the cytosol in intact cells (left panel, n=6) and into the buffer in isolated mitochondria (right panel, n=4). The amount of mtDNA was measured by qPCR. (**B**) SIRT3-depleted cells show enhanced expression of the *Ifnb1* gene in the basal state, as determined by qPCR (n=48). (**C**) SIRT3-depleted cells show enhanced expression of the *Ifnb1* gene following a 4-hour transfection with poly(I:C) and poly(dA:dT). (**D**) SIRT3-depleted cells exhibit elevated expression of multiple ISGs, including *Ifih1* (n=13), *Eif2ak2* (n=14), *Mx1* (n=14), *Ifitm1* (n=14), *Isg15* (n=21) and *Ifi44* (n=17), as assessed by qPCR.

In J774A.1 macrophages with *Sirt3* knockdown (**Fig. S1A**), interferon β mRNA levels were elevated at baseline (**Fig. 1B**) and following transfection with polyinosinic-polycytidylic acid (poly(I:C)) or poly(deoxyadenylic-deoxythymidylic) acid (poly(dA:dT)), synthetic analogs of viral RNA and DNA, respectively (**Fig. 1C**). These nucleic acids stimulate type I interferon responses through the RIG-I/MDA5-MAVS-IRF3 or the cGAS-STING-IRF3 axis respectively.

Consistent with this, *Sirt3* knockdown promoted expression of ISGs (Fig. 1D), including Interferon Induced with <u>H</u>elicase C Domain <u>1</u> (*Ifih1*, which codes for the RNA sensor MDA5), <u>E</u>ukaryotic Translation Initiation <u>F</u>actor <u>2 A</u>lpha <u>K</u>inase <u>2</u> (*Eif2ak2*, also called Protein kinase RNA-activated or protein kinase R [PKR]), Interferon-induced GTP-binding protein Mx1 (*Mx1*), Interferon-<u>s</u>timulated <u>g</u>ene <u>15</u> (*Isg15*), Interferon-Induced 44 (*Ifi44*) and Interferon-Induced <u>T</u>etratricopeptide repeat 1 (*Ifit1*). Given that PKR and MDA5 are key cytosolic RNA sensors, their elevated expression in Sirt3-depleted cells likely enhances the cellular responsiveness to dsRNA such as poly(I:C) (**Fig. 1D**), resulting in a more robust type I interferon response and amplified induction of multiple ISGs.

These findings were confirmed in human THP-1 macrophages, where transient SIRT3 knockdown amplified secretion of INF-β, with no change in TNF-α (**Fig. S1A,B**). Similar results were observed using CRISPR/Cas9-mediated *Sirt3* knockout in RAW 264.7 macrophages (**Fig. S1A,C**) and in primary bone marrow–derived macrophages (BMDMs) from mice with myeloid-specific *Sirt3* deletion (**Fig. S1A,D**). In all cellular models tested, SIRT3 deficiency enhanced IFN-β levels, demonstrating that SIRT3 broadly restrains type I interferon signaling.

### Type I interferons act as the paracrine signal released by Sirt3-depleted cells

Type I interferons function as both autocrine and paracrine mediators. We investigated whether elevated expression of ISGs in *Sirt3*-deficient cells might result from the increased release of IFN-β or other soluble mediators into the culture medium. To test this, control J774A.1 cells were incubated with conditioned medium from *Sirt3* knockdown cultures for 24 hours. This treatment induced robust expression of ISGs in control cells (**Fig. 2A**), indicating the presence of a paracrine soluble factor.

**Figure 2.**
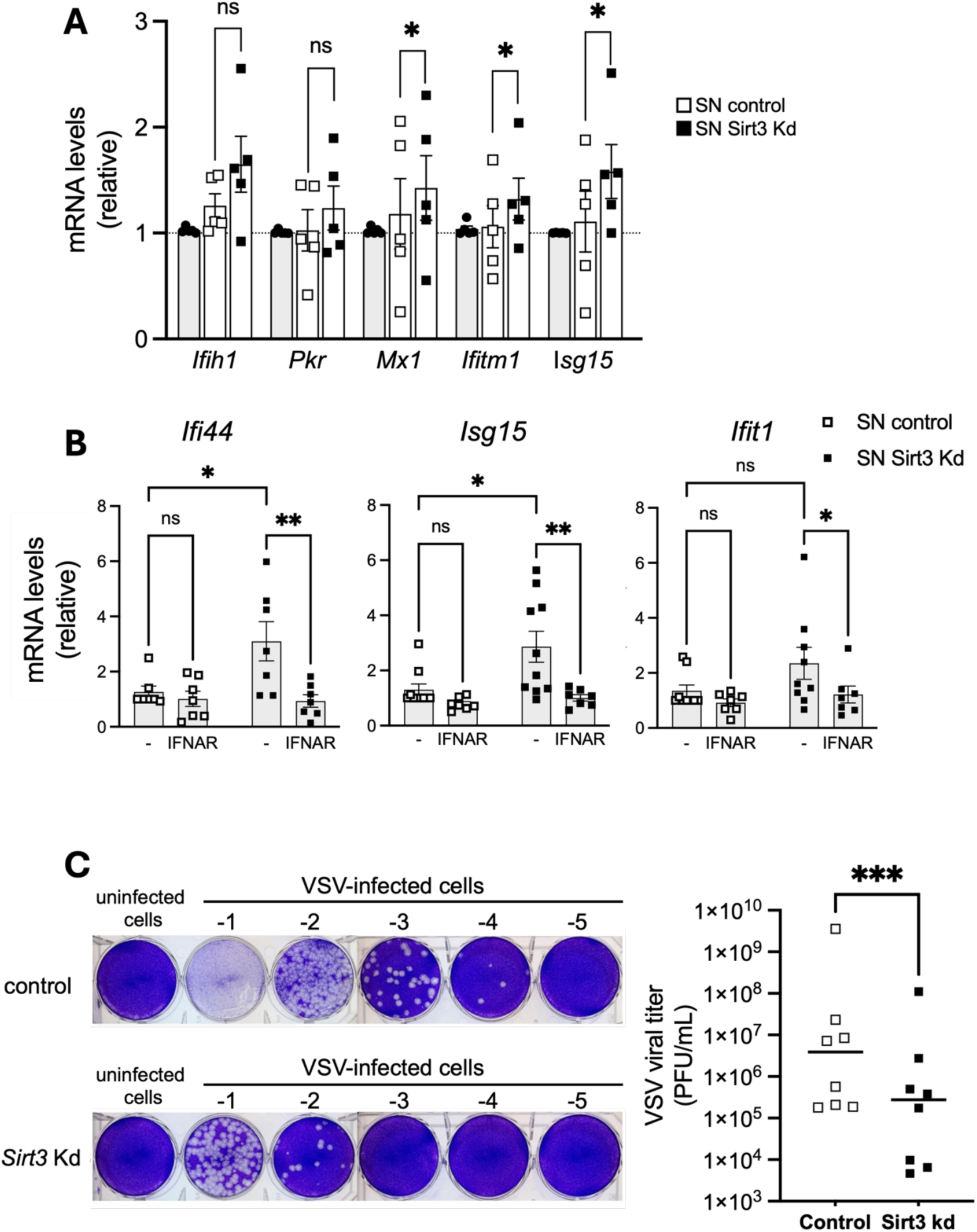
SIRT3 depletion induces the release of Type I interferons, which act as paracrine signals to enhance antiviral signaling. (**A**). Control J774A.1 cells were incubated for 24 hours with fresh medium (circles), their own supernatant (open squares) or supernatants obtained from SIRT3-depleted cells (closed squares). The mRNA levels of multiple ISGs were measured by qPCR, showing that conditioned medium from SIRT3 knockdown cells induced elevated ISG expression in control cells (n=5). (**B**) Control J774A.1 cells were incubated with supernatants from control or SIRT3-depleted cells in the presence or absence of blocking antibodies targeting the interferon-α/β receptor (IFNAR). ISG expression was determined by qPCR (n=7-10). (**C**) Viral resistance assays were performed in control and SIRT3-depleted J774A.1 macrophages infected with vesicular stomatitis virus (VSV) for 1 hour. Virus-containing supernatants were collected 24 hours post-infection and serially diluted (10-fold dilutions, 10⁻¹ to 10⁻⁵) for viral titration in BHK-21 cells. Representative images of BHK-21 cell monolayers stained with crystal violet after plaque formation are shown on the left. Viral titers, expressed as plaque-forming units per milliliter (PFU/mL), are quantified on the right (n=8).

The addition of neutralizing antibodies against the interferon-α/β receptor (IFNAR) abolished ISG induction (**Fig. 2B**), confirming that type I interferons mediate the response. These results demonstrate that IFN-β secreted by *Sirt3*-deficient cells acts in a paracrine manner, priming neighboring cells for enhanced antiviral and interferon-stimulated responses.

### SIRT3 restrains functional type I interferon signaling

To determine whether the increased type I interferon production associated with *Sirt3* knockdown was functionally active, we assessed its ability to induce an antiviral state. J774A.1 cells were infected with vesicular stomatitis virus (VSV, Indiana isolate) at MOI of 0.01. Viral supernatants were collected 24 hours post-infection, and viral titers were measured in BHK-21 cells. As shown in **Fig. 2C**, Sirt3 knockdown cells exhibited a ∼30-fold reduction in viral production compared with control cells. This reduction in viral replication is consistent with enhanced type I interferon signaling and ISG expression, demonstrating that the increased IFN-I response in *Sirt3*-deficient cells is biologically active and capable of inducing a robust antiviral state.

Thus, the VSV assay provides a functional readout of the heightened type I interferon response resulting from SIRT3 deficiency. Rather than reflecting an isolated increase in interferon production, these findings demonstrate that loss of SIRT3 leads to functionally amplified IFN-I signaling, providing a basis for examining its broader consequences for immune homeostasis and autoimmune inflammation.

### Type I interferon release by SIRT3-depleted cells requires intact mtDNA

To determine whether mtDNA contributes to the heightened IFN-β production in Sirt3-depleted cells, which display increased, mtDNA leak into the cytosol, we generated mtDNA-deficient cells (referred to as rho^0^ or ρ^0^ cells) by chronically treating them with 100 ng/ml ethidium bromide (EtBr), which inhibits mtDNA replication and depletes all 13 mitochondrially encoded proteins (**Fig. 3A**). Although ρ^0^ cells remain viable, they are deficient in oxidative phosphorylation (**Fig. 3A**) and require supplementation with uridine and pyruvate. Uridine is essential due to impaired dihydroorotate dehydrogenase activity in de novo pyrimidine synthesis, an enzyme localized in the inner mitochondrial membrane and dependent on the respiratory chain, while pyruvate supplementation supports aspartate synthesis, a process largely reliant on mitochondrial function (Birsoy et al., 2015; Sullivan et al., 2015).

**Figure 3.**
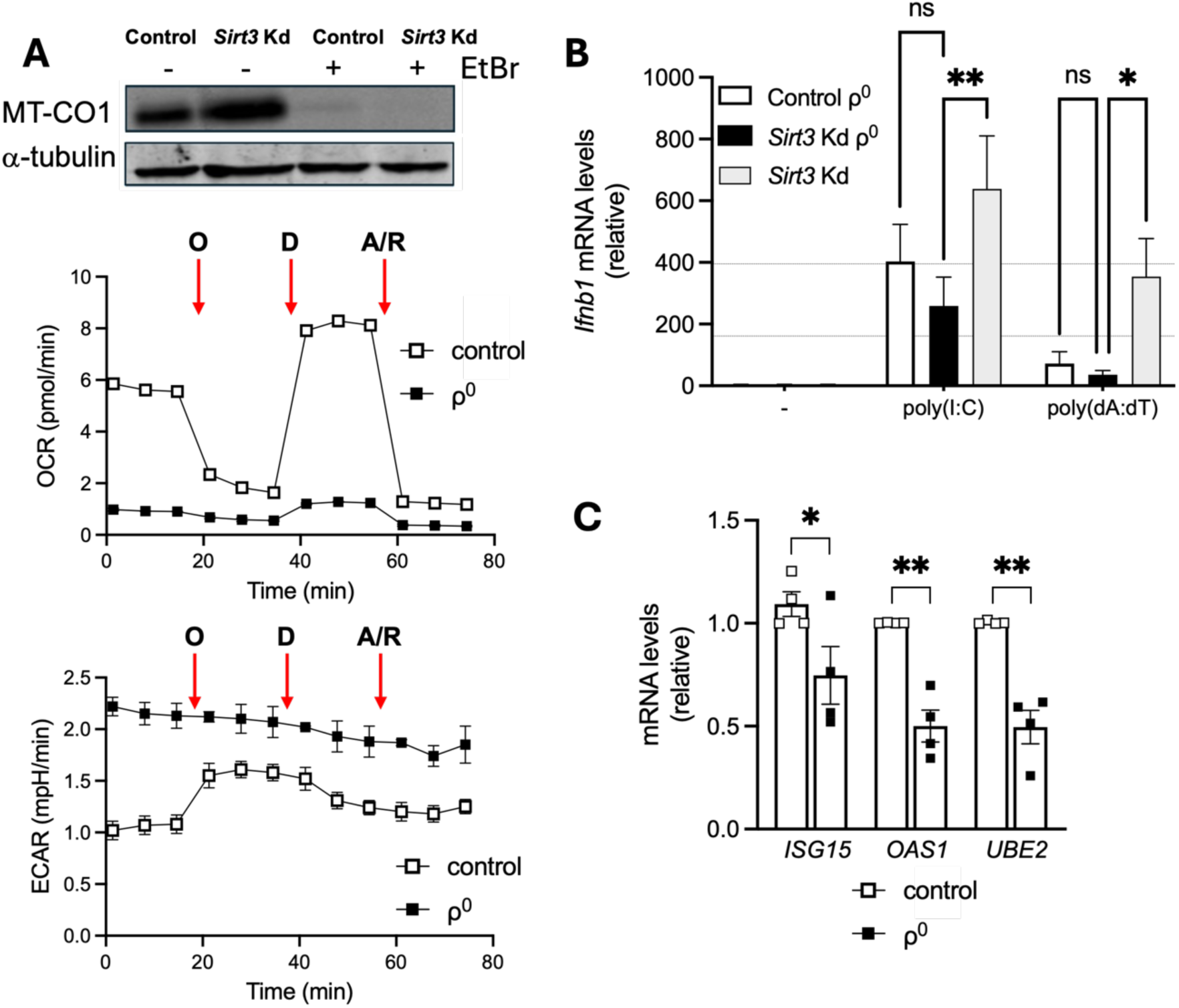
mtDNA is required for the enhanced type I interferon response in *Sirt3*-deficient cells. (**A**). Generation and characterization of mtDNA-depleted (ρ⁰) cells following chronic treatment with ethidium bromide (EtBr, 100 ng/mL) for 2 weeks. ρ⁰ cells were maintained in medium supplemented with uridine and pyruvate to support cell viability. The upper panel shows a representative western blot of MT-CO1, a mitochondrially encoded subunit of respiratory complex IV, with α-tubulin as a loading control. The middle panel shows oxygen consumption rate (OCR) measurements obtained using the Seahorse Mito Stress Test, demonstrating impaired mitochondrial respiration in ρ⁰ cells. The lower panel shows extracellular acidification rate (ECAR) from the same experiment, revealing increased basal glycolytic activity in ρ⁰ cells and the absence of the oligomycin-induced increase observed in WT cells. Seahorse data are from one representative experiment performed in triplicate. (**B**) *Ifnb1* gene expression in control and Sirt3-depleted ρ⁰ cells (n=10). Unlike mtDNA-containing cells, *Sirt3* knockdown did not increase *Ifnb1* expression in ρ⁰ cells. (**C**) ρ⁰ THP-1 cells exhibit decreased expression of multiple ISGs, including *ISG15, OAS1* and *UBE2*, compared to control THP-1 cells as assessed by qPCR (n=4).

In ρ⁰ cells, *Sirt3* knockdown failed to increase expression of the IFN-β gene (**Fig. 3B**). Similarly, ISG induction (*ISG15*, EIF2AK2, 2’-5’-oligoadenylate synthetase 1 (*OAS1*) and Ubiquitin-conjugating enzyme E2 L6 (*UBE2L6*)) was blunted in THP-1 ρ⁰ cells (**Fig. 3C**). Collectively, these results demonstrate that intact mtDNA is essential for the exaggerated type I interferon response observed in *Sirt3*-deficient cells.

### SIRT3-depleted cells display constitutive activation of the cGAS-STING pathway

Because mtDNA can activate cGAS–STING signaling, we examined pathway activation in SIRT3-depleted cells. Basal phosphorylation of STING and TBK1 was increased in J774A.1 and BMDM lacking *Sirt3* (**Fig. 4A**). Inhibition of cGAS with RU.521 reduced STING and TBK1 phoshorylation (**Fig. 4B**), and decreased IFN-β secretion in *Sirt3* knockdown cells (**Fig. 4C**), confirming that cytosolic mtDNA drives interferon signaling through the cGAS–STING axis in SIRT3-deficient cells.

**Figure 4.**
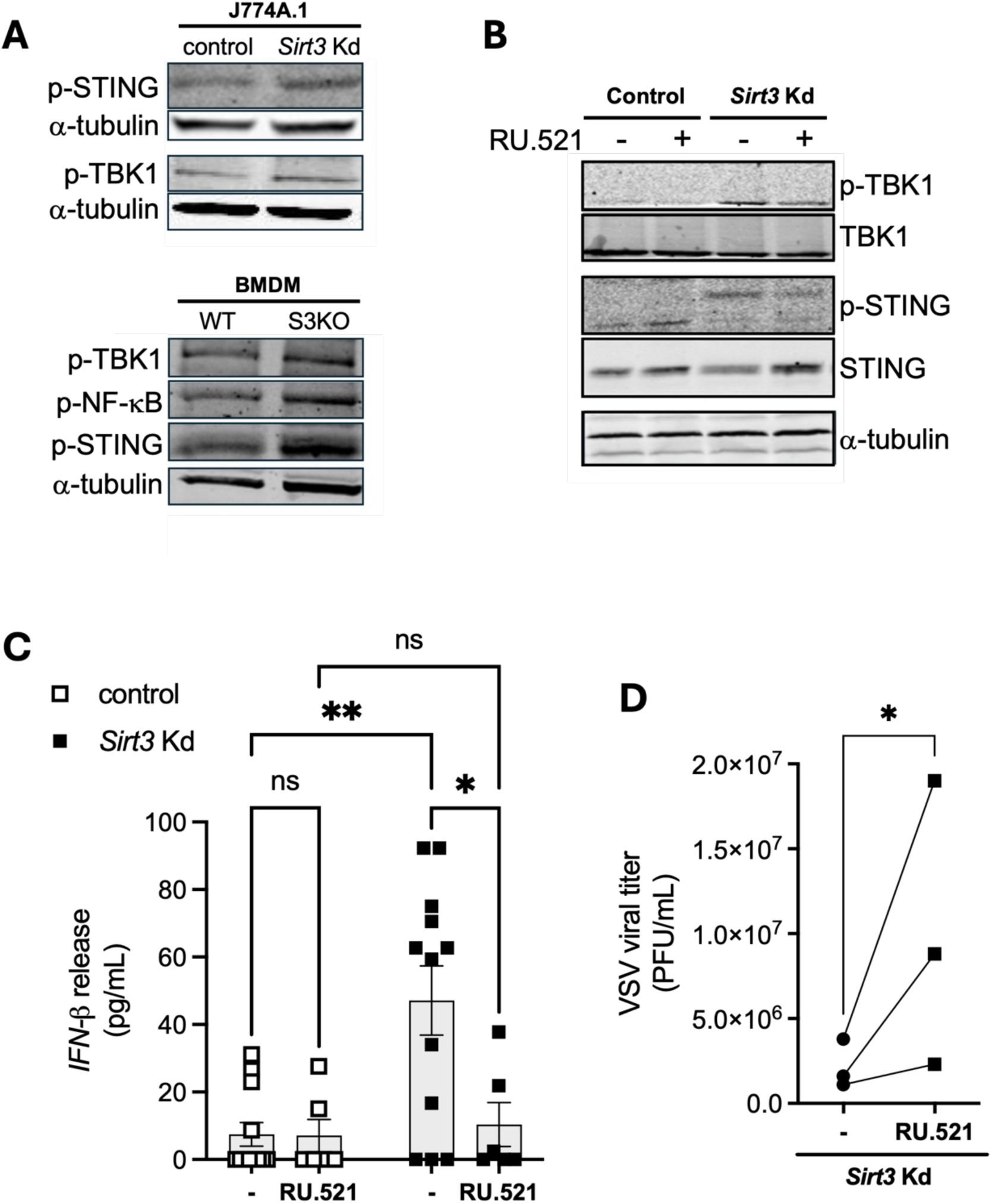
SIRT3 deficiency promotes cGAS-STING pathway activation and IFN-β production. (A) Basal phosphorylation of STING and TBK1 in control and SIRT3-depleted J774A.1 cells and in wild-type and *Sirt3*-knockout primary BMDMs, as assessed by immunoblotting. Increased phosphorylation of both signaling proteins indicates enhanced basal cGAS-STING pathway activity in the absence of SIRT3. (B) Effect of cGAS inhibition on STING and TBK1 phosphorylation in SIRT3-depleted J774A.1 cells following treatment with the murine cGAS inhibitor RU.521 (2 µg/mL, 24 h). (**C**) Effect of RU.521 treatment (2 µg/mL, 24 h) on IFN-β release by SIRT3-depleted cells (n=6–10). (**D**) Effect of RU.521 treatment (2 µg/mL, 24 h) on viral titers following VSV infection of SIRT3-depleted cells (n=3).

To determine whether the enhanced antiviral resistance observed in *Sirt3* knockdown J774A.1 cells is mediated by the cGAS-STING pathway, its signaling was inhibited. Sirt3-depleted cells were incubated with 20 µg/ml of the specific cGAS inhibitor RU.521 for 24 hours prior to infection. After this pre-treatment, cells were infected with VSV at an MOI of 0.01, and viral titers were quantified in BHK-21 cells (**Fig. 4D**). *Sirt3* knockdown cells pretreated with RU.521 displayed higher viral titers compared to untreated cells.

### mPTP-dependent release of oxidized mtDNA in SIRT3-deficient mitochondria

Regarding the mechanism of mtDNA release, we hypothesized that its efflux could occur through the mitochondrial permeability transition pore (mPTP), a multiprotein complex spanning the inner and outer mitochondrial membranes that opens in response to mitochondrial matrix Ca²⁺ accumulation and oxidative stress (Bernardi et al., 2023). To test this, Ca²⁺ uptake assays were performed in isolated mitochondria from control and Sirt3-depleted J774A.1 cells. Calcium Green was used to monitor extramitochondrial Ca²⁺, and sequential Ca²⁺ pulses were applied to progressively load mitochondria and induce mPTP opening. Mitochondrial Ca²⁺ uptake is reflected by a decrease in fluorescence, whereas mPTP opening results in Ca²⁺ release and a corresponding increase in fluorescence.

Following several Ca²⁺ additions, mitochondria from both control and Sirt3-depleted cells displayed a fluorescence increase consistent with mPTP opening (**Fig. 5A**). This response was abolished by cyclosporin A (CsA), an inhibitor of cyclophilin D (CypD), the main regulatory component of the mPTP (Baines et al., 2005), confirming that the observed Ca²⁺ release was mediated by mPTP opening (**Fig. 5B**). Exposure to ROS generated by the H_2_O_2_/Fe²⁺ system (García et al., 2005) accelerated mPTP opening in both control and Sirt3-depleted mitochondria, indicating that oxidative stress sensitizes the pore to opening, and this response was delayed by CsA (**Fig. 5C** and data not shown).

**Figure 5.**
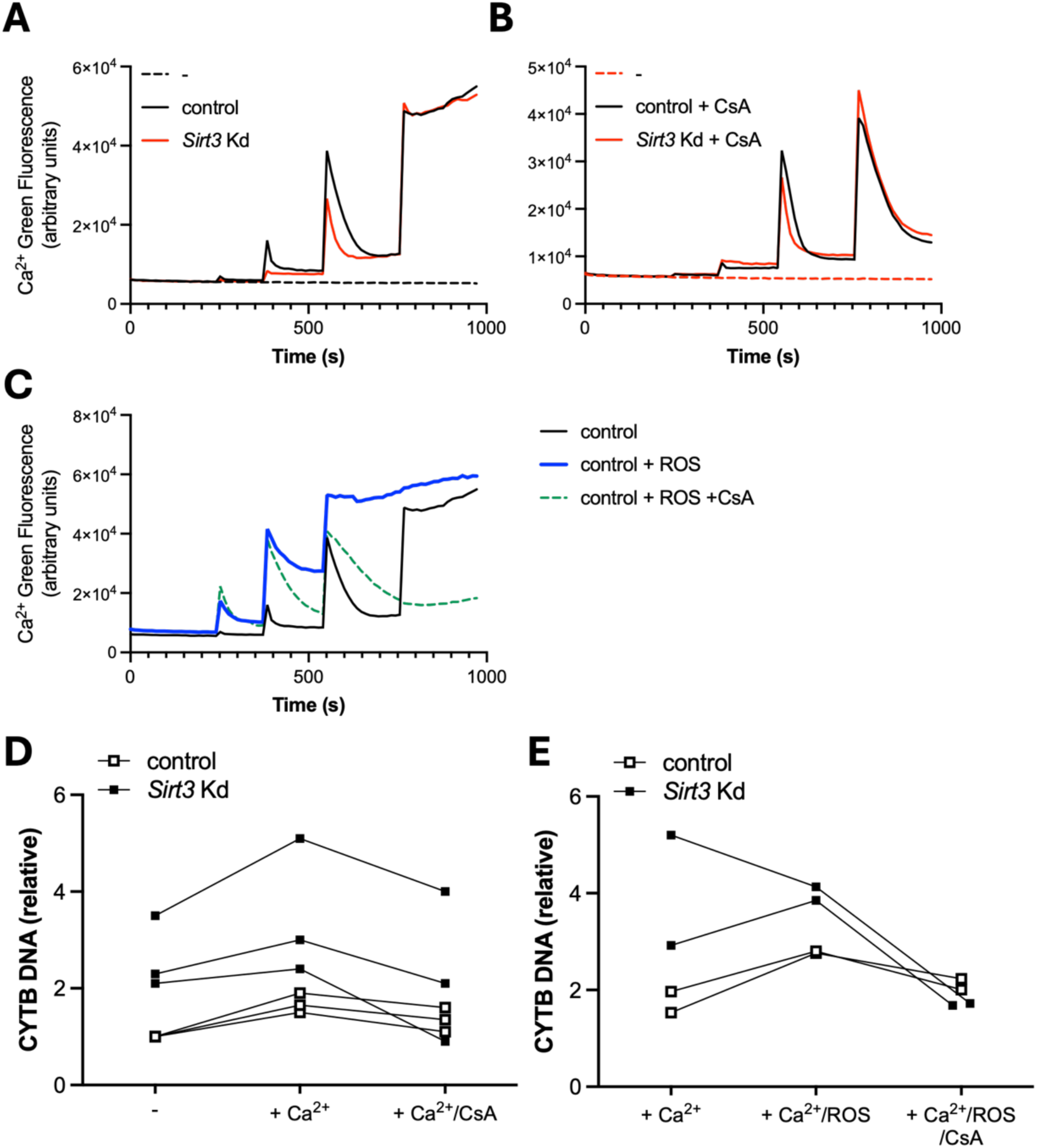
mPTP-dependent release of mtDNA in SIRT3-deficient mitochondria. (**A**–**C**) Mitochondrial permeability transition pore (mPTP) opening was assessed in isolated mitochondria from control and Sirt3-depleted J774A.1 cells by monitoring extramitochondrial Ca²⁺ using Calcium Green fluorescence. Sequential Ca²⁺ pulses were applied to induce mitochondrial Ca²⁺ loading and mPTP opening. (**A**) Representative fluorescence traces showing Ca²⁺ uptake and subsequent release following three sequential Ca²⁺ additions. (**B**) Effect of cyclosporin A (CsA) on Ca²⁺-induced mPTP opening. (**C**) Effect of oxidative stress generated by the H₂O₂/Fe²⁺ system on mPTP opening. (**D**) Quantification of mtDNA released into the incubation buffer under basal conditions or following Ca²⁺ stimulation, in the presence or absence of CsA. (**E**) Quantification of mtDNA released into the incubation buffer following Ca²⁺ stimulation, following Ca²⁺ stimulation in the presence of H₂O₂/Fe²⁺-induced oxidative stress and following Ca²⁺ stimulation in the presence of H₂O₂/Fe²⁺-induced oxidative stress and CsA. Data show that SIRT3 deficiency increases basal mtDNA release, whereas Ca²⁺-induced and ROS-enhanced mtDNA release are sensitive to CsA.

In parallel, mtDNA released into the incubation buffer was quantified under the same experimental conditions. In the absence of Ca²⁺ stimulation, mitochondria isolated from Sirt3-depleted cells exhibited significantly higher basal mtDNA release than control mitochondria (**Fig. 5D** and **Fig. 1A**). Ca²⁺ addition significantly enhanced mtDNA release in control mitochondria, whereas *Sirt3*-depleted mitochondria exhibited high basal release that did not significantly increase further upon Ca²⁺ stimulation.

Ca²⁺-induced mtDNA release was blocked by CsA, demonstrating that this process is mediated by the mPTP (**Fig. 5D**). ROS exposure enhanced mtDNA release in control mitochondria to levels comparable to those observed in Sirt3-depleted mitochondria, whereas the already elevated mtDNA release in Sirt3-depleted mitochondria in the presence of Ca²⁺ was not further increased by additional ROS (**Fig. 5E**). CsA inhibited mtDNA release in every experimental condition tested (**Fig. 5E**).

Collectively, these findings identify the mPTP as the route through which mtDNA is released from mitochondria. Although SIRT3 deficiency did not noticeably alter Ca²⁺-induced mPTP opening (**Fig. 5A**), it resulted in constitutively elevated basal mtDNA release through the pore (**Fig. 5D**). Together with our previous findings that SIRT3-deficient cells accumulate mitochondrial ROS (Traba et al., 2017), these results suggest that qualitative changes in mtDNA, rather than enhanced mPTP opening, underlie its increased basal escape from SIRT3-deficient mitochondria.

### mtDNA is oxidized and fragmented in the absence of SIRT3

SIRT3 is a mitochondrial deacetylase that plays a vital role in regulating oxidative stress by activating superoxide dismutase (SOD2), which converts superoxide radicals into hydrogen peroxide. In SIRT3-depleted cells, reduced SOD2 activity leads to increased production of mtROS (Qiu et al., 2010; Tao et al., 2010; Traba et al., 2017), which might activate innate immune pathways, including the cGAS-STING signaling cascade.

To determine the potential detrimental effect of elevated ROS on mtDNA, the status of mtDNA was analyzed in control and SIRT3-deficient cells. mtDNA oxidation was assessed by immunocytochemistry using an anti-8-oxo-dG antibody that specifically detects the oxidative lesion 7,8-dihydro-8-oxo-2ʹ-deoxyguanosine (8-oxo-dG). Analyses were performed under basal conditions and after 4 hours of LPS stimulation (**Fig. 6A-B**). Under basal conditions, SIRT3-deficient cells exhibited a significant increase in oxidized mtDNA compared with control cells. This increase was further enhanced upon LPS treatment.

**Figure 6.**
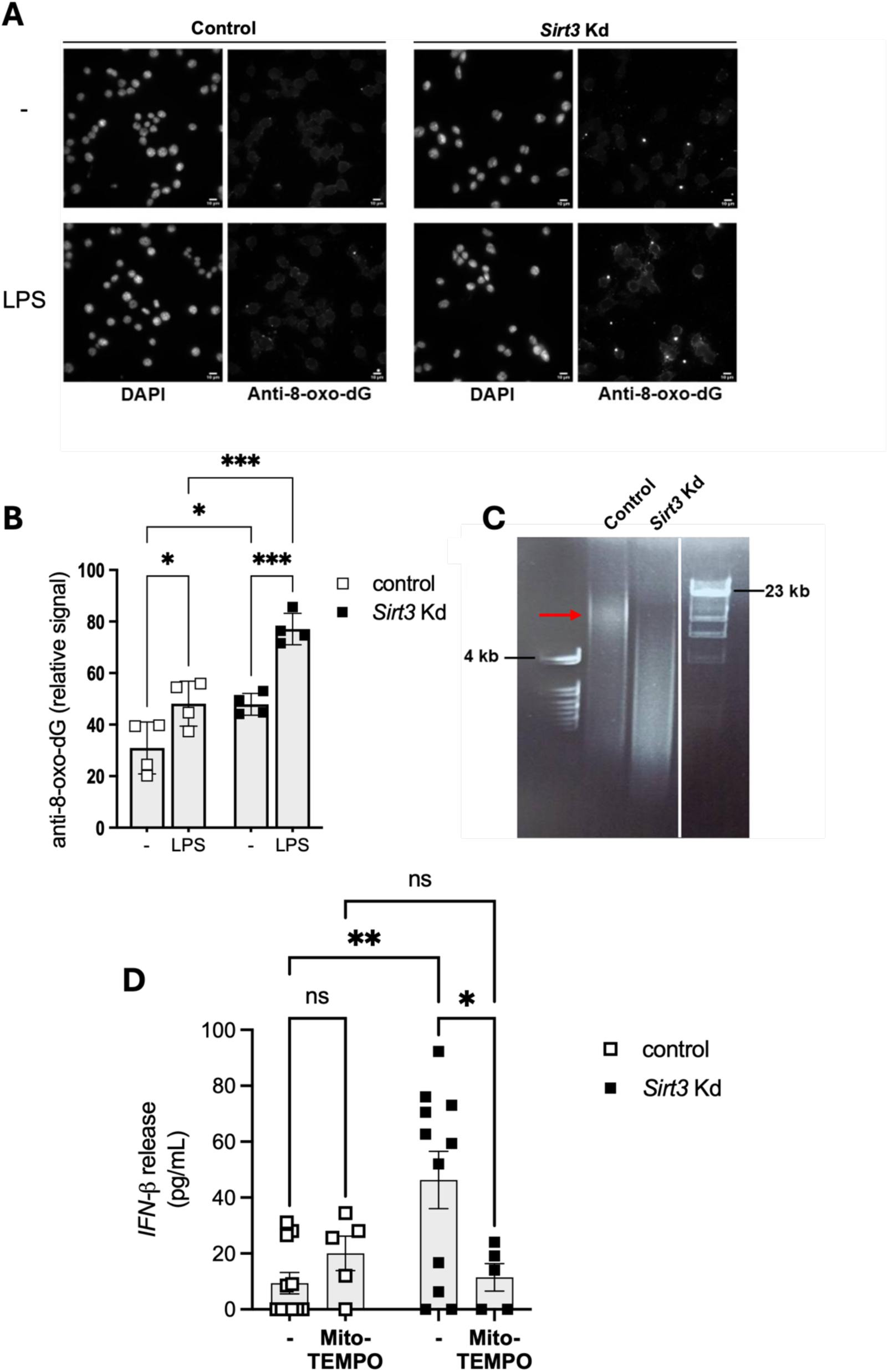
mtDNA is oxidized and fragmented in the absence of SIRT3. (**A**) Representative immunofluorescence images showing 8-oxo-dG staining in control and SIRT3-depleted J774A.1 cells under basal conditions (-) or following 4 h of LPS stimulation. (**B**) Quantification of 8-oxo-dG fluorescence intensity (n=4). (**C**) Agarose gel electrophoresis (1.2%) of mtDNA isolated from control and SIRT3-depleted J774A.1 cells. Ethidium bromide staining reveals intact mtDNA in control cells (red arrow) and increased mtDNA fragmentation in SIRT3-depleted cells. (**D**) FN-β release by control and Sirt3-knockdown J774A.1 cells following treatment with the mitochondrial superoxide scavenger Mito-TEMPO (10 µM, 24 h) (n=5–12).

We also examined the integrity of mtDNA from control and SIRT3-deficient cells. Mitochondrial DNA was extracted and resolved on agarose gels (**Fig. 6C**). Ethidium bromide staining revealed a higher number of DNA fragments smaller than 4,000 base pairs in SIRT3 depleted cells, whereas control cells predominantly displayed a band of approximately 16 kb, indicating increased mtDNA degradation in the absence of SIRT3. These results demonstrate that elevated ROS levels compromise mtDNA integrity in SIRT3-deficient cells.

Consistent with a role for mtROS in driving inflammation, treatment of *Sirt3* knockdown cells with Mito-TEMPO, a mitochondria-targeted superoxide scavenger, for 24 hours, significantly reduced the release of IFN-β in *Sirt3* knockdown cells (**Fig. 6D**). These findings indicate that mtROS are critical mediators of cGAS–STING pathway activation in SIRT3-deficient cells.

### SIRT3 protects against STING–dependent inflammatory tissue damage

While transient type I interferon responses are essential for antiviral defense, persistent activation of the cGAS-STING/type I interferon axis is increasingly recognized as a driver of chronic inflammatory and autoimmune disorders. We therefore investigated whether the enhanced cGAS-STING signaling observed in SIRT3-deficient macrophages also increases susceptibility to STING-dependent inflammatory tissue injury in vivo.

Chlorhexidine gluconate (CHX) induces STING-dependent peritoneal fibrosis in mice (Marchant et al., 2025). To determine whether SIRT3 protects against this form of inflammatory tissue injury, we evaluated peritoneal injury following daily intraperitoneal injections of 0.1% CHX for 10 days (**Fig. 7A**).

**Figure 7.**
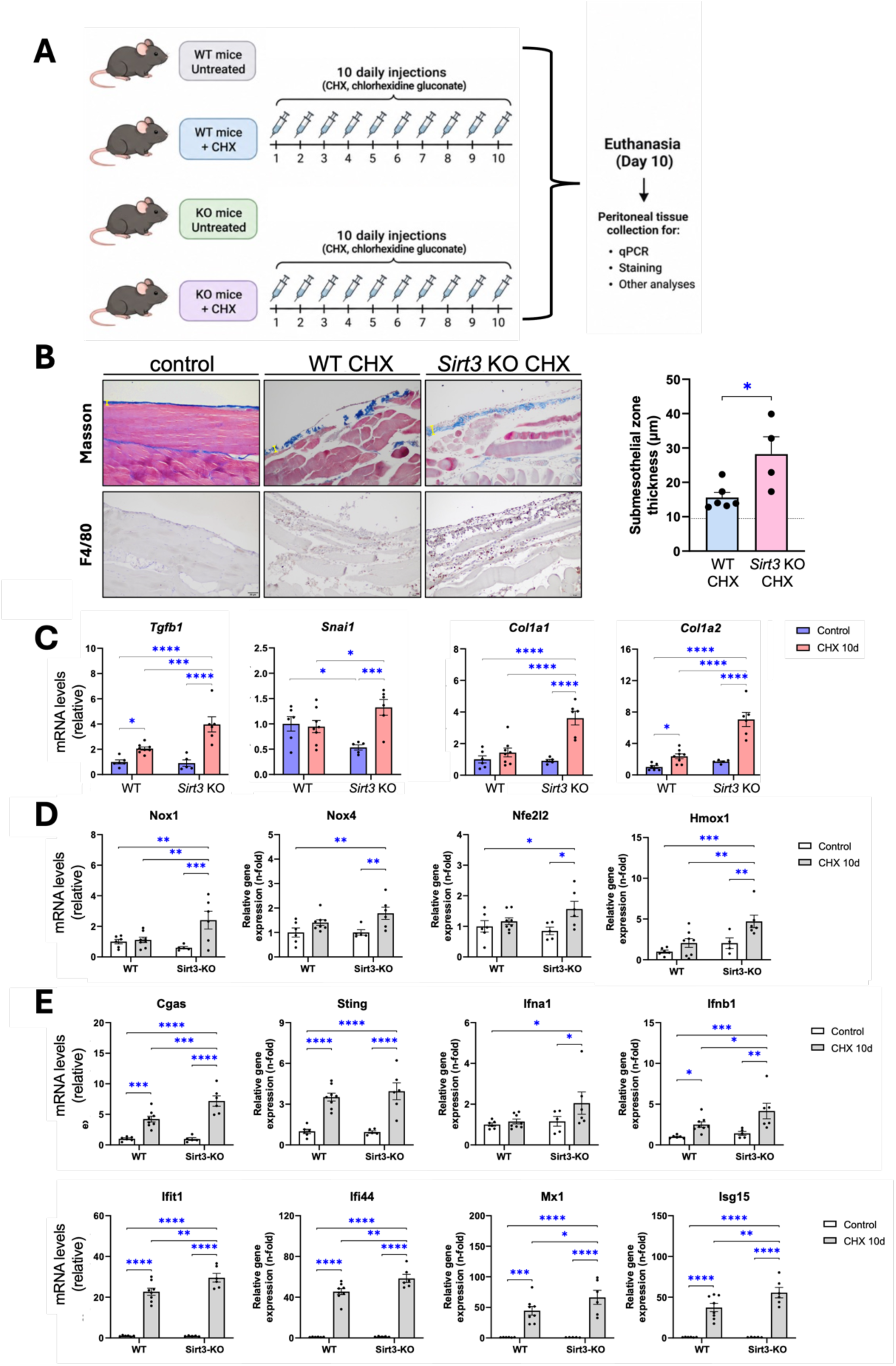
SIRT3 deficiency is associated with increased peritoneal remodeling and macrophage infiltration following CHX treatment. (**A**) Experimental design. WT and Sirt3-KO mice were left untreated or treated daily with chlorhexidine gluconate (CHX) for 10 days. At the end of the treatment period, mice were euthanized and peritoneal tissue was collected for histological and molecular analyses (n = 6 untreated WT, 5 untreated *Sirt3* KO, 8 CHX-treated WT mice and 6 CHX-treated *Sirt3* KO mice). (**B**) Representative Masson’s trichrome staining (upper panels) and F4/80 immunostaining (lower panels) of peritoneal tissue. Masson’s trichrome staining was used to assess the thickness of the submesothelial zone, while F4/80 staining was used to evaluate macrophage infiltration. Increased macrophage infiltration was observed in CHX-treated *Sirt3* KO mice. The histogram on the right shows the quantification of submesothelial zone thickness. (**C**) Relative mRNA expression of the fibrosis-associated genes *Tgfb1* (encoding the transforming growth factor beta 1), *Snai1*, *Col1a1* (encoding the α1 chain of type I collagen), and *Col1a2* (encoding the α1 chain of type II collagen) in peritoneal tissue from untreated and CHX-treated WT and Sirt3-KO mice, as determined by qPCR. (**D**) Relative mRNA expression of genes associated with oxidative stress and antioxidant defense, including *Nox1* and *Nox4* (encoding NADPH oxidases), *Nfe2l2* (encoding the antioxidant transcription factor NRF2), and *Hmox1* (encoding heme oxygenase-1), in peritoneal tissue, as determined by qPCR. (**E**) Relative mRNA expression of genes associated with the cGAS-STING pathway and type I interferon responses, including *Mb21d1* (cGAS), *Tmem173* (STING), *Ifna1* (IFN-α), *Ifnb1*, and the ISGs *Ifit1*, *Ifi44*, *Mx1*, and *Isg15*, in peritoneal tissue, as determined by qPCR.

Histological analysis revealed that CHX exposure led to a significant increase in the thickness of the submesothelial zone and in macrophage infiltration in myeloid-specific *Sirt3* knockout mice compared with wildtype controls (**Fig. 7B**), as visualized in Masson’s trichrome and F4/80-stained sections, respectively.

At the molecular level, genes associated with mesothelial-to-mesenchymal transition and fibrosis (*Tgfb1, Snai1, Col1a1, and Col1a2*), as well as oxidative stress–related genes (*Nox1, Nox4, Nfe2l2, and Hmox1*), showed a progressive upregulation in response to prolonged CHX exposure, particularly in *Sirt3*-deficient mice (**Fig. 7C–D**). These changes were accompanied by an increase in the expression of cGAS-STING and type I-IFN response components (**Fig. 7E**).

Together, these findings demonstrate that SIRT3 protects against STING-dependent inflammatory tissue injury and fibrotic remodeling *in vivo,* extending our *in vitro* observations to a physiologically relevant disease model.

### SIRT3 downregulation is associated with autoimmune disease in humans

Given the pathological consequences of SIRT3 deficiency in mice, we next investigated whether reduced SIRT3 expression is also associated with human autoimmune disease. To address this question, we reanalyzed publicly available transcriptomic datasets from previous studies (Bertier et al., 2012; Kang et al., 2017) using the GEO2R analysis tool. We first examined glomerular tissue from patients with lupus nephritis (GEO dataset GSE32591). UMAP analysis was used to visualize the distribution of the samples (**Fig. 8A**), while individual expression profiles revealed an approximately 30% reduction in SIRT3 transcript levels compared with healthy controls (**Fig. 8B**). Differential expression analysis further identified changes in the expression of SIRT3 and several interferon-stimulated genes (ISGs) (**Fig. 8C**). In particular, the expression of SIRT3 was reduced, whereas that of the ISGs MX1, ISG15, and IFI44 was increased in lupus nephritis samples (**Fig. 8D**). Sample characteristics and probe-level statistical analyses are provided in **Tables S1, S3**.

**Figure 8.**
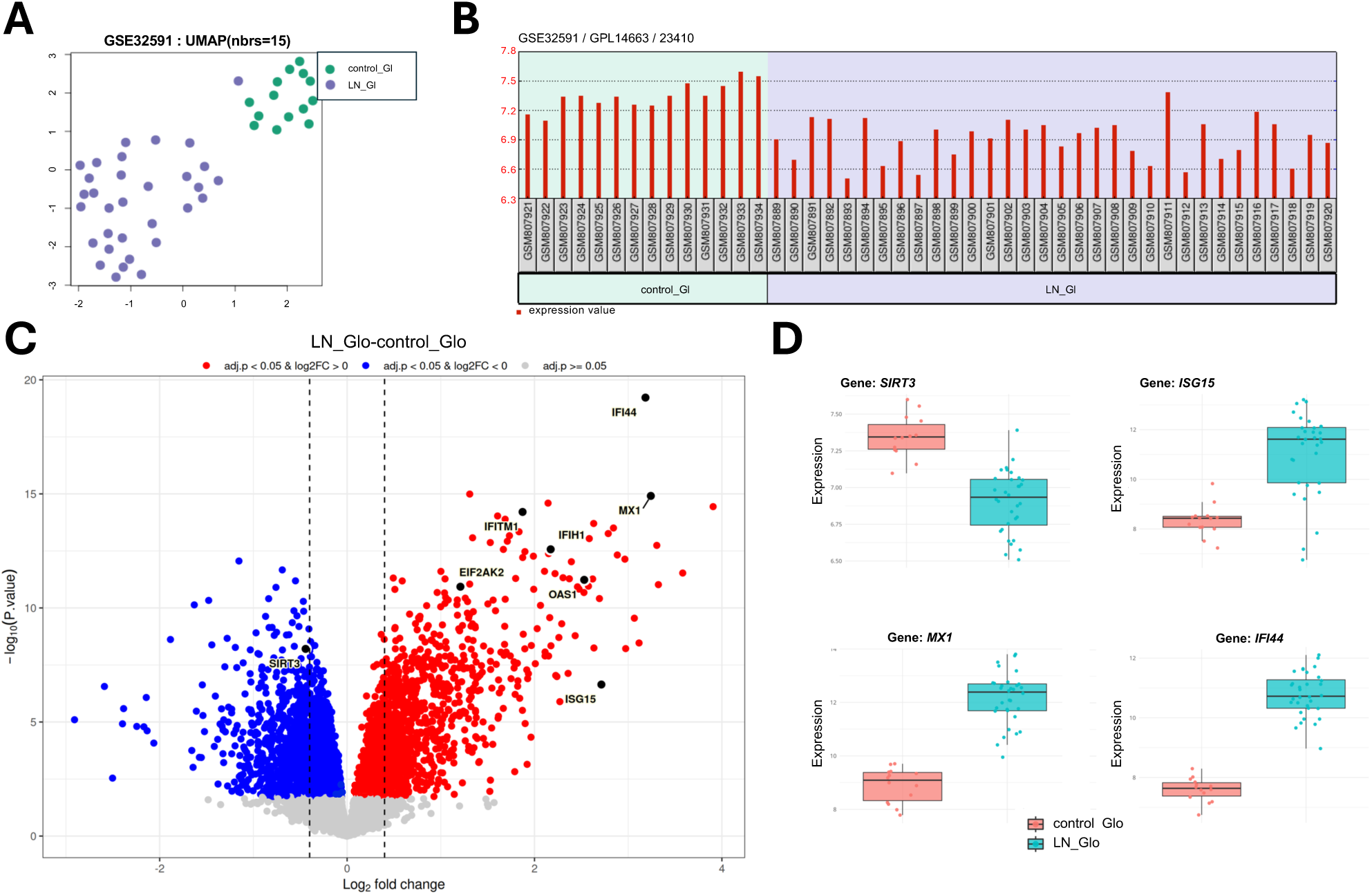
Reduced SIRT3 expression and increased ISG expression in glomerular tissue from patients with lupus nephritis. (**A**) Uniform Manifold Approximation and Projection (UMAP) plot showing the distribution of glomerular samples from patients with lupus nephritis and healthy controls (GEO dataset GSE32591). (**B**) Individual SIRT3 expression levels in glomerular samples from each participant. (**C**) Volcano plots showing differential gene expression of SIRT3 and selected ISGs between lupus nephritis patients and healthy controls. (**D**) Box plots showing the expression levels of SIRT3 and the ISGs MX1, ISG15, and IFI44 in glomerular samples from both groups. Transcriptomic data were reanalyzed using GEO2R. Sample characteristics and probe-level statistical analyses are provided in **Tables S1, S3**.

We next assessed whether these findings extended to other renal compartments in the same study and to other autoimmune conditions. A similar analysis of tubulointerstitial tissue from the same lupus nephritis cohort revealed a comparable expression pattern (**Supplementary Fig. S2**). Furthermore, analysis of macrophages from patients with rheumatoid arthritis (GEO dataset GSE97779) showed an approximately 46% reduction in SIRT3 expression, together with increased expression of ISGs (**Supplementary Fig. S3**). Sample characteristics and probe-level statistical analyses are provided in **Tables S2, S4**. The expression profiles of additional ISGs in both datasets are presented in **Supplementary Fig. S4**.

Together, these findings show that reduced SIRT3 expression is associated with an enhanced interferon-related transcriptional signature across distinct tissues and autoimmune conditions. In combination with our murine data showing increased type I interferon production in the absence of SIRT3, these observations support a conserved association between SIRT3 deficiency and enhanced interferon responses across species and suggest that the mechanism identified in our experimental models may also be relevant to human inflammatory disease.

## DISCUSSION

SIRT3 plays a pivotal role in limiting inflammation across various contexts. In a murine model of sepsis-induced acute kidney injury (AKI) following caecal ligation and puncture, SIRT3 exhibited protective effects by reducing ROS production and inhibiting the NLRP3 inflammasome (Zhao et al., 2016). Another study demonstrated that SIRT3 regulates the acetylation of autophagy-related proteins, with SIRT3-deficient macrophages displaying impaired autophagy and accelerated NLRP3 inflammasome activation (Liu et al., 2018). In a model of hyperglycemic intracerebral hemorrhage, SIRT3 improved brain edema and reduced neuronal apoptosis by enhancing resistance to oxidative stress and modulating the SIRT3/SOD2/NLRP3 pathway (Zheng et al., 2018). SIRT3 activation, induced by agents like honokiol or melatonin, has shown therapeutic potential in conditions such as atherosclerosis and AKI, primarily by promoting mitophagy and inhibiting the NLRP3 inflammasome (Ma et al., 2018; Deng et al., 2023).

Furthermore, SIRT3 activation has been reported to reduce NLRP3 activation and mitigate pathogenic effects in a *Pseudomonas aeruginosa*-induced pneumonia model (Maurice et al., 2022). More recently, SIRT3 has been shown to inhibit the cGAS-STING pathway in both a mouse model of silicosis and hepatic ischemia–reperfusion injury, suggesting broader anti-inflammatory effects of SIRT3 (Kong et al., 2024; Zhou et al., 2024). The present study expands on the role of SIRT3 by demonstrating its critical function in controlling the release of mitochondrial DNA (mtDNA) and the downstream activation of innate immune pathways, specifically type I interferon signaling.

Our findings reveal that SIRT3 deficiency in macrophages promotes a basal increase in mtDNA leakage, even in resting cells, which serves as a potent trigger for type I interferon responses through the cGAS-STING pathway. This basal mtDNA leak appears to fine-tune the antiviral defenses of the cell, priming cells for an amplified response upon viral or synthetic nucleic acid challenges. The elevated release of IFN-β in SIRT3 knockdown cells underscores the regulatory effect of SIRT3 in balancing immune responses, as the excess interferon also acts in a paracrine manner, enhancing antiviral defenses in neighboring cells.

An important aspect of this response concerns the mechanism underlying mtDNA release. Our data do not support enhanced mPTP opening in SIRT3-deficient mitochondria. Instead, they suggest that increased mtDNA release may reflect changes in the mitochondrial DNA substrate rather than an increase in pore opening. Elevated mitochondrial ROS in SIRT3-deficient cells is associated with increased mtDNA oxidation and fragmentation, potentially facilitating its passage through transient mPTP openings. In this regard, García et al. (2005) reported that mtDNA released through the mPTP consisted of fragments smaller than 1,000 bp, suggesting that fragmentation may facilitate the release of mtDNA through this pathway, given the substantially larger size of the intact mitochondrial genome (approximately 16.5 kb). Although the precise contribution of mtDNA fragmentation to its release remains to be established, these observations provide a possible explanation for the increased cytosolic mtDNA detected in SIRT3-deficient cells without evidence of enhanced mPTP opening.

In line with previous studies demonstrating the anti-inflammatory effects of SIRT3 across various inflammatory and oxidative stress-related contexts, our results suggest that the regulatory impact of SIRT3 extends to modulating antiviral responses. The observed upregulation of interferon-stimulated genes (ISGs) and reduced viral replication in SIRT3-deficient cells provide strong evidence that SIRT3 restricts the antiviral state. Furthermore, experiments with ρ^0^ cells lacking mtDNA confirmed that mtDNA is a critical mediator in the exaggerated type I interferon response observed in SIRT3-deficient cells, as loss of mtDNA in these cells abrogated the increased IFN-β and ISG expression. These results highlight mtDNA as a key signal driving the enhanced immune response in the absence of SIRT3.

Our study also identifies mtROS as a pivotal factor linking SIRT3 deficiency to cGAS-STING pathway activation. With reduced SOD2 activity in SIRT3-depleted cells, elevated mtROS likely drives the release of mtDNA and subsequent immune signaling. The reduction in type I interferon and ISG expression upon treatment with the mitochondria-targeted redox-active antioxidant Mito-TEMPO supports this mechanism, underscoring the role of oxidative stress in activating the cGAS-STING pathway when SIRT3 is absent. This finding aligns with previous studies in which the role of SIRT3 in reducing ROS has been shown to mitigate inflammatory pathways such as NLRP3 inflammasome activation.

Together, these findings suggest that SIRT3 not only limits inflammatory pathways like the NLRP3 inflammasome but also acts as a gatekeeper for antiviral responses by regulating mtDNA release and ROS production. The ability of SIRT3 to inhibit the cGAS-STING pathway, as reported in recent studies in models of silicosis and hepatic ischemia–reperfusion injury (Kong et al., 2024; Zhou et al., 2024), further supports its broad role in modulating innate immunity. By controlling mtDNA release and ROS, SIRT3 prevents inappropriate activation of type I interferon responses, which, if unchecked, may contribute to pathological inflammation and autoimmune conditions. Thus, SIRT3 emerges as a critical regulator that balances the need of the immune system to defend against pathogens while avoiding excessive immune activation.

Interestingly, a recent study reported that SIRT3 may enhance antiviral immunity through the deacetylation of mitochondrial antiviral-signaling protein (MAVS) (Liu et al., 2024), providing a potentially contrasting view of SIRT3 function. MAVS is localized to the outer mitochondrial membrane, whereas SIRT3 is predominantly located in the mitochondrial matrix. This difference in subcellular localization raises questions regarding the accessibility of MAVS acetylation sites to SIRT3 and the mechanism underlying the reported interaction. More broadly, these findings suggest that SIRT3 may exert distinct effects on antiviral signaling depending on the pathways involved. While the present study identifies an inhibitory role for SIRT3 in mtDNA-dependent cGAS–STING signaling, its reported effects on MAVS raise the possibility that SIRT3 differentially regulates distinct branches of innate antiviral immunity.

Our in vivo studies further support the protective role of SIRT3 in limiting inflammation and tissue damage. In the chlorhexidine gluconate-induced model of peritoneal fibrosis, SIRT3 deficiency was associated with increased macrophage infiltration, enhanced expression of ISGs, and more pronounced fibrotic remodeling. Previous work by Marchant et al. (2026) established the involvement of STING signaling in this model of peritoneal fibrosis. In this context, our findings extend these observations by showing that SIRT3 deficiency exacerbates the inflammatory response associated with STING activation. The increased expression of interferon-related genes, together with macrophage accumulation and fibrotic changes, supports a role for mitochondrial homeostasis in controlling the progression of STING-associated tissue inflammation and remodeling. These results also highlight the potential consequences of sustained mtDNA-driven innate immune activation beyond cultured macrophages.

Finally, reanalysis of publicly available gene-expression datasets revealed reduced SIRT3 expression in glomerular samples from patients with lupus nephritis and in macrophages from patients with rheumatoid arthritis (Berthier et al., 2012; Kang et al., 2017). These observations indicate that reduced SIRT3 expression is also associated with human inflammatory conditions and support the relevance of investigating its role in the regulation of innate immune responses. Although these datasets do not establish a causal relationship between SIRT3 downregulation and interferon signaling in patients, the findings are consistent with the possibility that altered SIRT3 expression may contribute to inflammatory processes in human disease.

Collectively, our findings identify SIRT3 as an important regulator of mitochondrial integrity and innate immune signaling. By limiting mitochondrial oxidative stress and mtDNA release, SIRT3 restrains basal cGAS–STING activation and type I IFN production, thereby influencing both cell-autonomous and paracrine antiviral responses. These findings have potential therapeutic implications, as modulation of SIRT3 activity could help regulate interferon responses in different disease contexts. While reducing SIRT3 activity might enhance antiviral defenses under certain conditions, increasing its activity could be beneficial in pathological settings characterized by excessive innate immune activation and tissue inflammation. However, the reported effects of SIRT3 on distinct antiviral signaling pathways, including MAVS and cGAS–STING, suggest that the consequences of targeting SIRT3 will depend on the biological context. Further investigation of these pathways may help define how SIRT3 modulation could be used to fine-tune innate immune response.

## MATERIALS AND METHODS

### Cell culture

J774A.1 mouse macrophages (ATCC) were maintained in Dulbecco’s Modified Eagle Medium (DMEM) supplemented with 10% heat-inactivated FBS. Stable control and *Sirt3* knockdown J774A.1 cells were generated previously (Traba et al., 2017) using puromycin-selectable lentiviral shRNA constructs (Sigma).

THP-1 human monocyte cells (ATCC) were cultured in Roswell Park Memorial Institute 1640 (RPMI 1640) medium supplemented with 25 mM HEPES and 10% heat-inactivated fetal bovine serum (FBS). To induce differentiation into macrophages, cells were treated with 5 ng/ml phorbol 12-myristate 13-acetate (PMA) for 48 h. Transient siRNA transfection was performed by electroporation 18 hours prior to differentiation, as previously described (Traba et al., 2015).

RAW 264.7 macrophages (ATCC) were cultured in DMEM supplemented with 10% heat-inactivated FBS. A single guide RNA (sgRNA) targeting the mouse *Sirt3* gene (sequence: 5ʹ-ATGTCACTCACTACTTCCTG-3ʹ) was designed using the Benchling online tool (benchling.com). Forward and reverse oligonucleotides were synthesized with appropriate overhangs for cloning into the BpiI-digested pX459 vector (Fw: 5ʹ-CACCG-[20 nt]-3ʹ; Rv: 5ʹ-AAAC-[20 nt reverse complement]-C-3ʹ). RAW 264.7 cells were transfected with either pX459-Sirt3 sgRNA, or the empty pX459 vector, and subjected to puromycin selection for 3 days to enrich successfully transfected cell and facilitate the generation of *Sirt3* knockout lines.

Primary bone marrow–derived macrophages (BMDMs) were prepared using standard protocols (Anton & Traba, 2022). Briefly, bone marrow cells isolated from mouse femurs and tibiae were seeded on sterile culture dishes and incubated for 7 days in DMEM supplemented with 25 mM HEPES, 10% heat-inactivated FBS, and 20% L929 cell-conditioned medium.

Baby Hamster Kidney-21 (BHK-21) cells were cultured in DMEM supplemented with 5% heat-inactivated FBS and were used as host cells for viral titration assays.

### Mouse models

All mice used in this study were in a C57Bl/6 J genetic background. Cre-loxP-mediated recombination was employed to specifically delete SIRT3 in cells from myeloid origin. Homozygous floxed SIRT3 mice (SIRT3^fl/fl^) [B6.129(Cg)-Sirt3tm1.1Auw/J] were crossed with transgenic mice expressing Cre recombinase under the control of the *Lyz2* (lysozyme 2) promoter [B6.129P2-Lyz2tm1(cre)Ifo/J]. The resulting offspring included SIRT3^fl/fl^ LysM-Cre^−/−^ controls and SIRT3^fl/fl^ LysM-Cre^+/−^, hereafter referred to as SIRT3 KO.

Peritoneal damage was induced in mice by daily intraperitoneal (i.p.) injections of 0.1% chlorhexidine gluconate (CHX) diluted in 0.9% NaCl saline solution, as previously described by Marchant et al., (2025). CHX was administered at a volume of 10 mL/kg body weight for 10 consecutive days.

All animal procedures complied with the European Directive 86/609/EEC for the protection of animals used for scientific purposes, and were approved by the CSIC Ethics Committee, the Animal Welfare Ethics Committee of the IIS-FJD, and the Comunidad de Madrid (PROEX 220.8/23 and 242.2/21).

### Generation of mtDNA-deficient (ρ^0^) cells

Mitochondrial DNA–deficient (ρ⁰) cells were generated by culturing cells in medium supplemented with ethidium bromide (EtBr, 100 ng/mL) for 2 weeks, together with sodium pyruvate (100 μg/mL) and uridine (50 μg/mL) to support cell viability in the absence of mitochondrial oxidative phosphorylation. Following EtBr treatment, cells were maintained in EtBr-free medium. The continued requirement for exogenous pyruvate and uridine was used as a functional indicator of mtDNA depletion and loss of mitochondrial respiratory capacity.

mtDNA depletion was confirmed by quantitative real-time PCR (qPCR) targeting mitochondrial-encoded genes and by immunoblotting for mitochondrially encoded proteins, including cytochrome c oxidase subunit I (MT-CO1; Abcam).

### Cell stimulation and cytokine assays

J774A.1, RAW 264.7, BMDMs, and THP-1-derived macrophages were transfected with poly(I:C) or poly(dA:dT) (InvivoGen) using Lipofectamine 3000 (Invitrogen), according to the manufacturer’s instructions. Four hours after transfection, culture supernatants were collected and centrifuged to remove cells and cellular debris. Clarified supernatants were stored at −80 °C until cytokine analysis. IFN-β and TNF-α concentrations were determined using ELISA kits (R&D Systems). In parallel, cells were harvested at the same time point for RNA and protein extraction, as described below.

### Immunoblot analysis

Cells were lysed in radioimmunoprecipitation assay (RIPA) buffer, and total protein extracts were separated by SDS-PAGE and transferred onto nitrocellulose membranes. Inmunodetection was performed using the Odyssey infrared imaging system (LI-COR Biosciences). The following primary antibodies were used: phospho-STING (72971, Cell Signaling Technology), STING (13647S, Cell Signaling Technology), phospho-TBK1 (5483S, Cell Signaling Technology), TBK1 (3504S, Cell Signaling Technology) SIRT3 (5490S, Cell Signaling Technology) and a-tubulin (T6199, Sigma),

To detect mitochondrially encoded proteins, an antibody against MT-CO1 (ab14705, Abcam) was used. For MT-CO1 immunoblotting, samples were not boiled before gel loading, according to the manufacturer’s instructions.

### Quantitative PCR analysis

Total RNA was extracted from cells using TRIzol reagent (Thermo Fisher Scientific), and cDNA was synthesized using a first-strand synthesis kit (Thermo Fisher Scientific), according to the manufacturer’s protocol. Quantitative real-time PCR (qPCR) was performed using QuantiTect Primer Assays (Qiagen) or custom-designed primers (Thermo Fisher Scientific, **Table S5**). Relative gene expression was calculated using the 2⁻ΔΔCt method, with Ct values normalized to the expression of reference (housekeeping) genes.

#### Cytosolic mtDNA quantification

Cytosolic mtDNA was quantified by qPCR as previously described (Anton and Traba, 2022), with modifications. Briefly, cells were treated with digitonin to selectively permeabilize the plasma membrane, allowing the release of cytosolic contents into the incubation medium. The medium was collected, and DNA was isolated using the DNeasy Blood and Tissue Kit (Qiagen). The copy number of the mitochondrial DNA-encoded cytochrome b (*Cytb*) was quantified by qPCR using the following custom-designed primers synthesized by Thermo Fisher Scientific: forward, 5ʹ-GCTTTCCACTTCATCTTACCATTTA-3ʹ; and reverse, 5ʹ-TGTTGGGTTGTTTGATCCTG-3ʹ.

#### Immunofluorescence detection of 8-oxodG

J774A.1 cells were seeded at low density onto coverslips and cultured at 37°C in a humidified atmosphere containing 5% CO₂. Cells were either left untreated or stimulated with LPS (100 ng/mL) for 4 h. Following PBS washing, cells were fixed with methanol for 5 min at room temperature and treated with RNase A (5 mg/mL; Sigma) in 5 mM Tris-HCl (pH 7.5) and 7.5 mM NaCl for 1 h at 37°C to remove RNA. Cells were subsequently washed with PBS containing 0.1% Triton X-100 (PBD), refixed with 4% paraformaldehyde (PFA), and washed again. DNA denaturation was achieved by incubation with 50 mM NaOH in 50% ethanol for 7 min at room temperature. After washing, nonspecific binding was blocked with PBS containing 0.2% Triton X-100 and 1% BSA for 45 min at room temperature.

Cells were incubated with anti-8-oxodeoxyguanosine antibody (8-oxodG; Santa Cruz Biotechnology; 1:100) for 1 h at room temperature, followed by Alexa Fluor 488-conjugated donkey anti-mouse IgG (Thermo Fisher Scientific; 1:400) for 1 h. Nuclei were counterstained with DAPI (1:1000). Coverslips were mounted using Mowiol-DABCO and allowed to dry overnight at 4°C in the dark.

Images were acquired using a Zeiss inverted microscope equipped with a CCD camera and either a 63×/1.2 water-immersion objective or a 10×/0.45 apochromatic objective.

### Viral infections

J774A.1 macrophages were infected with vesicular stomatitis virus (VSV, Indiana isolate) at a multiplicity of infection (MOI) between 0.001 and 0.1. Cytopathic effects (CPE) were monitored microscopically 4–5 hours post-infection. Viral supernatants were collected at 24 hours post-infection, and viral titers were determined by plaque assay in BHK-21 cells. For plaque visualization, BHK-21 monolayers were fixed with 2% paraformaldehyde and stained with 2% crystal violet

### Mitochondrial calcium uptake experiments

Mitochondrial Ca²⁺ uptake and mPTP opening were assessed essentially as previously described (Traba et al., 2012). Briefly, mitochondrial Ca^2+^ uptake was monitored using the Ca^2+^-sensitive fluorescent probe Calcium-Green 5N (0.1 μM; excitation, 506 nm; emission, 532 nm) in isolated mitochondria, which were resuspended in intracellular (IC) buffer (125 mM KCl, 2 mM potassium phosphate, 1 mM MgCl_2_, and 20 mM Hepes, pH 7.4).

Fluorescence was recorded using an OPTIMA FLUOstar plate reader equipped with temperature control and continuous stirring. Measurements were performed at 30°C in the presence of succinate (5 mM) and rotenone (1 μM) to support mitochondrial respiration. Following a 3–5 min equilibration period, sequential CaCl₂ pulses were added to progressively load mitochondria with Ca²⁺ until mPTP opening was detected, as indicated by the release of Ca²⁺ into the extramitochondrial medium and the consequent increase in fluorescence. Where indicated, cyclosporin A (CsA, 5 μM) was added to inhibit mPTP opening. To promote ROS generation through the Fenton reaction, FeSO₄ (400 μM) and H₂O₂ (3 mM) were added as indicated (García et al., 2005).

### Chlorhexidine gluconate-induced peritoneal damage mouse model

The chlorhexidine gluconate (CHX)-induced peritoneal injury model was performed essentially as previously described (Marchant et al., 2025). Briefly, peritoneal injury was induced in WT and SIRT3-deficient (SIRT3-KO) male mice (9-12 weeks old) by daily intraperitoneal injections of 0.1% CHX dissolved in saline solution (0.9% NaCl) at a volume of 10 mL/kg of weight. Mice were either left untreated or injected daily with CHX for 10 days (WT Control, n = 7; WT CHX, n = 6; SIRT3-KO Control, n = 5; SIRT3-KO CHX, n = 7). At the experimental endpoint, mice were euthanized, peritoneal lavages were collected, and parietal peritoneal tissues were harvested for histological and molecular analyses.

Peritoneal tissues were fixed in 4% paraformaldehyde, paraffin-embedded, and sectioned at 4 μm for Massońs trichrome staining (Bio-Optica, Milano, Italy) or immunohistochemistry (IHC). For IHC, tissue sections were deparaffined and hydrated, and antigen retrieval was carried out using a sodium citrate buffer (10 mM. pH 6 or 9) on a PT Link system (DAKO, Glostrup, Denmark). Endogenous peroxidase blockade was done using 3% hydrogen peroxide (Millipore, Burlington, MA, USA) and protein blockade using a casein-blocking solution (Vector Laboratories, Newark, CA, USA). Then rat anti-F4/80 (1:50; MCA497, Bio-Rad, CA, USA) primary antibodies were incubated overnight at 4 °C, followed by incubation with biotinylated secondary antibodies (anti-rat, 1:200) and avidin-biotin complex (ABC, Vector Laboratories). Signal was detected using 3,3-diaminobenzidine (DAB) chromogen and substrate solution (Abcam, Cambridge, UK). Finally, slides were counterstained with hematoxylin (Merck, Darmstadt, Germany), dehydrated, and mounted with DPX (Merck).

Peritoneal membrane thickness was quantified by measuring the width of the submesothelial compact zone on Masson’s trichrome-stained sections using ImageJ (five measurements per field in 5–10 fields per mouse; ×200 magnification). F4/80-positive cells were quantified by counting positive cells in 5–10 randomly selected fields per mouse (×200 magnification).

For gene expression analysis, total RNA was isolated from frozen peritoneal tissue using TRItidy G (PanReac) according to the manufacturer’s instructions. cDNA was synthesized from 2 μg of total RNA using the High-Capacity cDNA Reverse Transcription Kit (Applied Biosystems). Quantitative PCR was performed using Premix Ex Taq (Takara) together with predesigned TaqMan assays for target and housekeeping genes (Supplementary Table S3) on 7500 Fast Real-Time PCR and QuantStudio 3 instruments (Applied Biosystems). Relative gene expression was calculated using the 2^−ΔΔCt method after normalization to housekeeping genes and expressed as fold change relative to control samples.

### Transcriptomic analysis of human autoimmune datasets

Publicly available transcriptomic datasets were analyzed to assess the association between SIRT3 expression and type I interferon–related gene signatures in human autoimmune diseases. The datasets GSE32591 (lupus nephritis, LN; Berthier et al., 2012) and GSE97779 (rheumatoid arthritis, RA; Kang et al., 2017) were retrieved from the Gene Expression Omnibus (GEO) database. Differential expression analyses were performed using GEO2R, and the resulting data were further processed using R (v4.3).

In the LN dataset, glomerular and tubulointerstitial samples from patients were analyzed separately against samples from healthy controls. In the RA dataset, synovial macrophages from patients were compared with control macrophages from healthy individuals. Differentially expressed genes were identified using the Benjamini–Hochberg method for multiple-testing correction, with an adjusted P value < 0.05 and an absolute log₂ fold change ≥ 0.4 as selection criteria. Particular attention was given to SIRT3 and a set of interferon-stimulated genes (ISGs), including ISG15, MX1, IFI44, IFIT1, and OAS1.

### Statistical analysis

Statistical analyses were performed using PraphPad Prism (version 11). Comparison between two groups were performed using paired or unpaired two-tailed y-tests as appropriate. One-way ANOVA was used when more than two groups were compared. Two-way ANOVA was applied when two independent variables were analyzed. A p-value of < 0.05 was considered statistically significant.

## ACKNOWLEDGEMENTS

We thank the laboratory of Jorgina Satrústegui, Araceli del Arco and Beatriz Pardo (CBM) for their continuous support and valuable discussions. We are also grateful to Irene García-Higuera (CBM) for providing the B6.129P2-Lyz2tm1(cre)Ifo/J mouse. Data analysis was carried out at the Biocomputational Analysis Core Facility (https://www.cbm.uam.es/index.php/facilities/biocomputational-analysis/) of the Centro de Biología Molecular Severo Ochoa (CSIC-UAM, Madrid, Spain), which receives institutional funding from both CSIC and UAM.

## CONFLICT OF INTEREST

None declared

**Supplementary Figure S1. SIRT3-depleted mouse and human cells exhibit enhanced type I interferon responses.** (**A**) Representative western blots showing SIRT3 protein levels and α-tubulin as a loading control in the four experimental models used in this study: J774A.1 macrophages stably expressing control or *Sirt3*-targeting shRNA (previously generated in Traba et al., 2017); THP-1-derived macrophages transiently transfected with control (siCT) or *SIRT3*-targeting siRNA; RAW264.7 macrophages with permanent *Sirt3* disruption generated using CRISPR/Cas9 (S3KO); and bone marrow-derived macrophages (BMDMs) from mice with myeloid-specific *Sirt3* deletion using LysM-Cre. (**B**) *SIRT3* was knocked down in THP-1-derived macrophages using siRNA and, following a 4-hour stimulation with poly(I) and poly(dA), IFN-β protein levels in the supernatant were measured by ELISA (n=5). (**C**) IFN-β protein levels in the supernatant of wild-type or *Sirt3*-knockout RAW264.7 cells were measured by ELISA (n=3). (**D**) IFN-β protein levels in the supernatant of BMDM obtained from wild-type or myeloid-specific *Sirt3*-knockout mice were measured by ELISA (n=8). (**E**) BMDMs obtained from myeloid-specific *Sirt3*-knockout mice exhibit elevated expression of *Ifnb* (n=5) and multiple ISGs, including *Ifi44* (n=4), *Isg15* (n=5), and *Ifit1* (n=5), as assessed by qPCR.

**Supplementary Figure S2. Reduced SIRT3 expression and increased ISG expression in tubulointerstitial tissue from patients with lupus nephritis.**

(**A**) Uniform Manifold Approximation and Projection (UMAP) plot showing the distribution of tubulointerstitial samples from patients with lupus nephritis (LN) and healthy controls (GEO dataset GSE32591). (**B**) Individual SIRT3 expression levels in tubulointerstitial samples from each participant. (**C**) Volcano plots showing differential gene expression between LN patients and healthy controls. (**D**) Box plots showing the expression levels of SIRT3 and the ISGs MX1, ISG15, and IFI44 in both groups. Transcriptomic data were reanalyzed using GEO2R. Sample characteristics and probe-level statistical analyses are provided in **Tables S1, S3**.

**Supplementary Figure S3. Reduced SIRT3 expression and increased ISG expression in macrophages from patients with rheumatoid arthritis.**

(**A**) Uniform Manifold Approximation and Projection (UMAP) plot showing the distribution of macrophage samples from patients with rheumatoid arthritis (RA) and healthy controls (GEO dataset GSE97779). (**B**) Individual expression levels of SIRT3 in macrophage samples from each participant, showing the results obtained with the three probes corresponding to SIRT3. (C) Volcano plots showing differential gene expression between RA patients and healthy controls. (D) Box plots showing the expression levels of SIRT3, MX1, ISG15, and IFI44 in both groups. Three probes were used to assess SIRT3 expression, two probes for IFI44, and one probe for each of the remaining genes. Transcriptomic data were reanalyzed using GEO2R. Sample characteristics and probe-level statistical analyses are provided in **Tables S2, S4**.

**Supplementary Figure S4. Expression of additional interferon-stimulated genes in human autoimmune disease datasets.**

Expression levels of additional interferon-stimulated genes (ISGs), not included in **Fig. 8** or **Supplementary Fig. S1** and **Supplementary Fig. S2**, in (**A**) glomerular and (**B**) tubulointerstitial samples from patients with lupus nephritis (LN) and healthy controls (GEO dataset GSE32591), and (**C**) macrophages from patients with rheumatoid arthritis (RA) and healthy controls (GEO dataset GSE97779). Transcriptomic data were reanalyzed using GEO2R.

**Table S1. GSE32591 (lupus nephritis) samples and related metadata.** GEO accession numbers, sample titles, source descriptions, disease status, patient identifiers, and tissue type (glomerular or tubulointerstitial) for all samples included in the transcriptomic dataset.

**Table S2. GSE97779 (rheumatoid arthritis) samples and related metadata.** GEO accession numbers, sample titles, source descriptions, cell status, subject disease status, subject identifiers, and tissue of origin for all control and RA macrophage samples included in the transcriptomic dataset.

**Table S3. Differential expression analysis of SIRT3 in glomerular and tubulointerstitial tissues from the lupus nephritis cohort (GSE32591).** Results of the GEO2R analysis showing the adjusted p-value (adj.P.Val), nominal p-value (P.Value), moderated t-statistic (t), B-statistic (B), and log fold change (logFC) for SIRT3 in glomerular and tubulointerstitial samples.

Table S4.**Probe-level differential expression analysis of SIRT3 in the rheumatoid arthritis cohort (GSE97779).** Results of the GEO2R analysis for the three probes corresponding to SIRT3, showing probe identifiers (Probe ID), adjusted p-values (adj.P.Val), nominal p-values (P.Value), moderated t-statistics (t), B-statistics (B), and log fold changes (logFC).

**Table S5.**
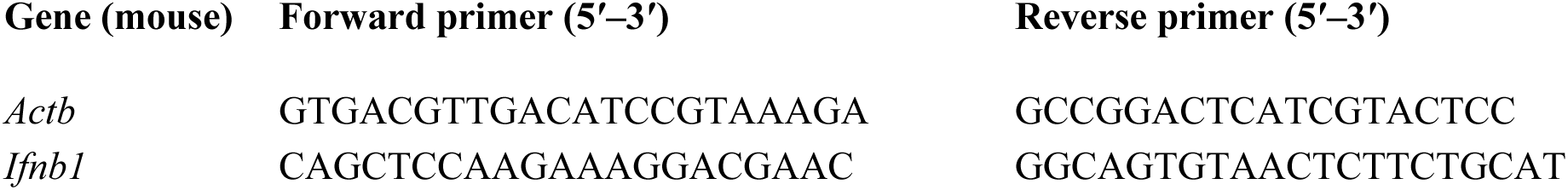
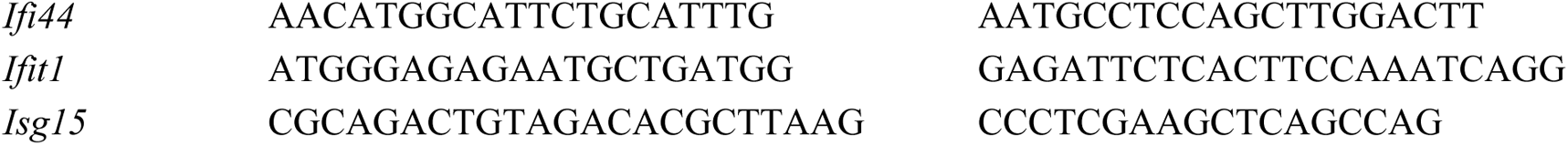
Sequences of custom-designed primers used for quantitative real-time PCR.

