## Supplemental Figures for "SIRT3 preserves mitochondrial DNA integrity to restrain cGAS–STING signaling and autoimmune inflammation"

**Figure S1**.

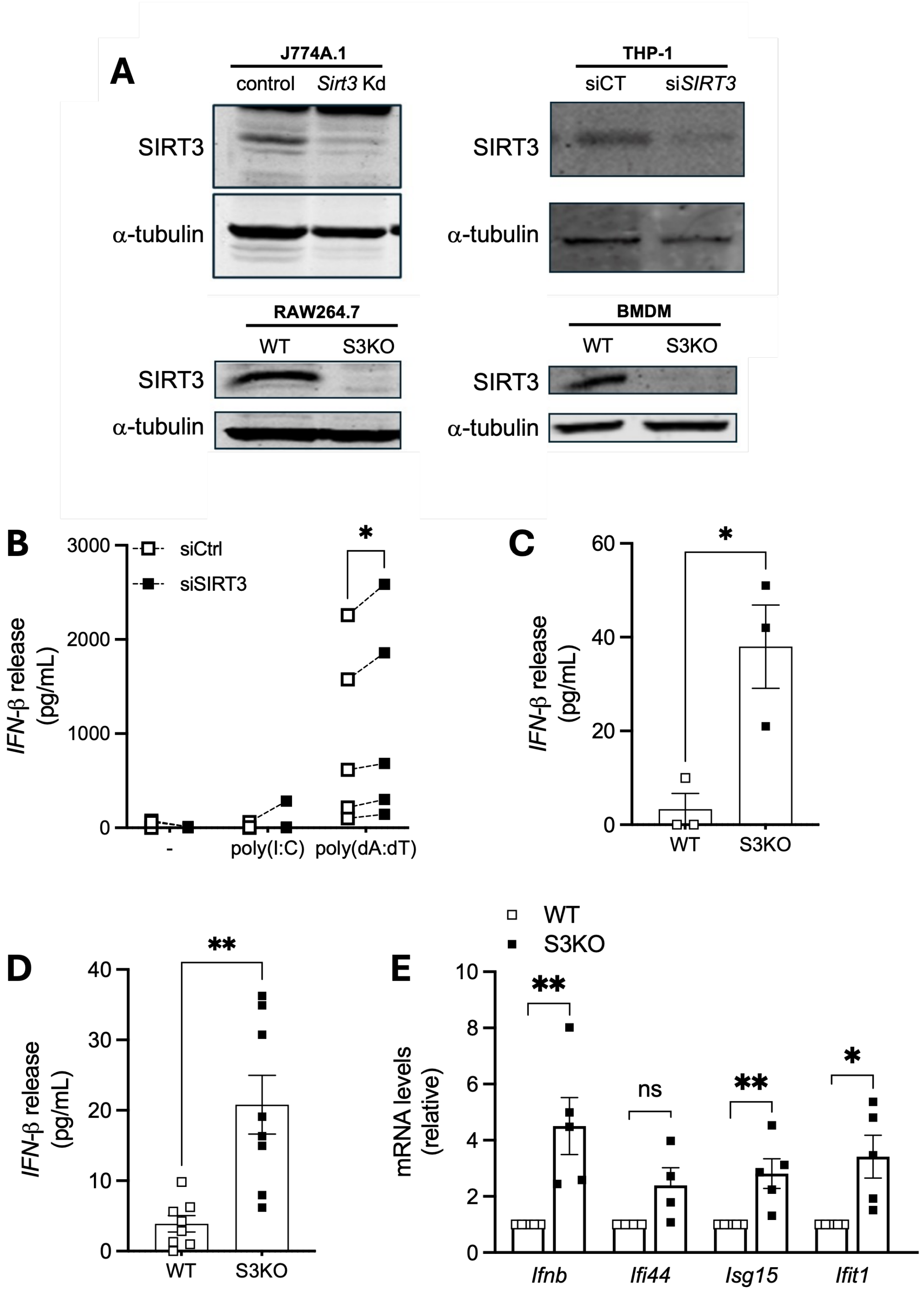
A
J774A.1
control
Sirt3 Kd
SIRT3
α-tubulin
THP-1
siCT
siSIRT3
RAW264.7
WT
S3KO
BMDM
B
siCtrl
siSIRT3
IFN-β release (pg/mL)
-
poly(I:C)
poly(dA:dT)
C
D
E
mRNA levels (relative)
ns
Ifnb
Ifi44
Isg15
Ifit1

**Figure S2**.

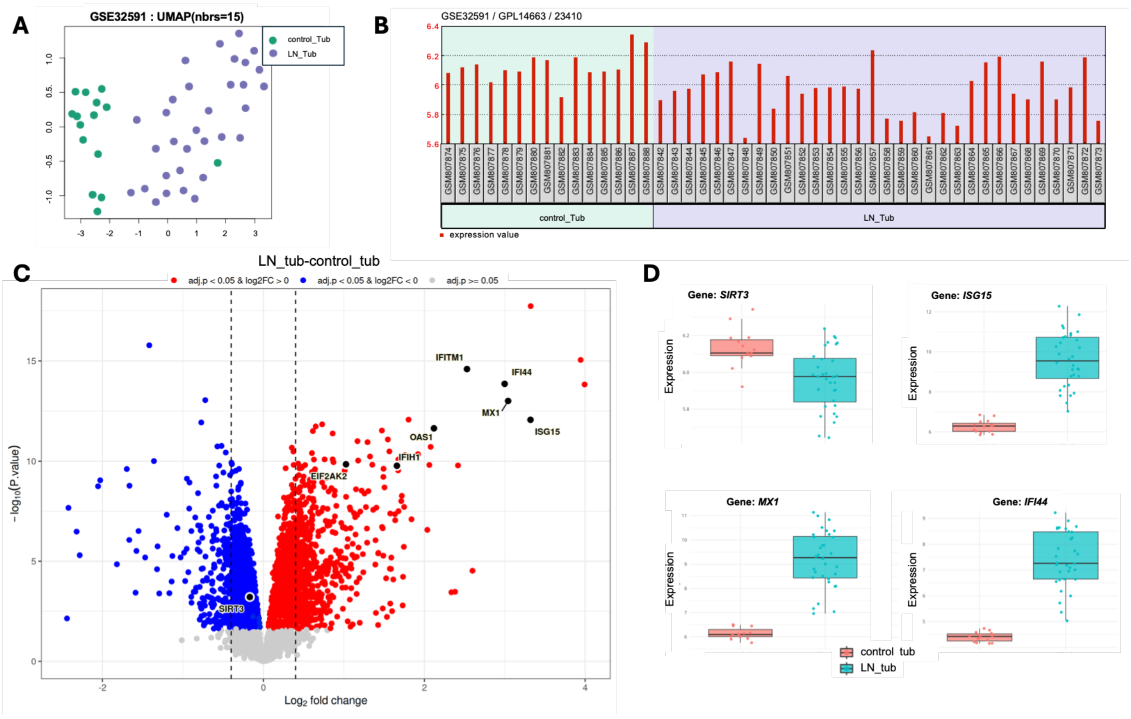
A
GSE32591 : UMAP(nbrs=15)
control_Tub
LN_Tub
B
GSE32591 / GPL14663 / 23410
control_Tub
LN_Tub
expression value
C
LN_tub-control_tub
adj.p < 0.05 & log2FC > 0
adj.p < 0.05 & log2FC < 0
adj.p >= 0.05
−log10(P.value)
Log2 fold change
IFITM1
IFI44
MX1
ISG15
OAS1
IFIH1
EIF2AK2
SIRT3
D
Gene: SIRT3
Gene: ISG15
Gene: MX1
Gene: IFI44
Expression
control_tub
LN_tub

### Figure S3.

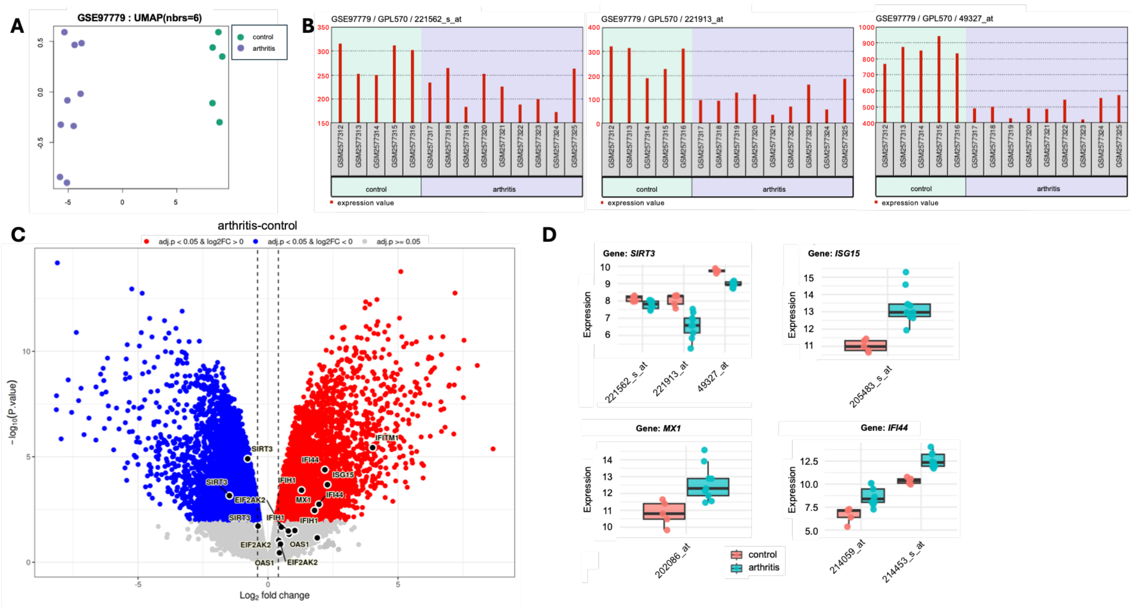
A
GSE97779 : UMAP(nbrs=6)
control
arthritis
B
GSE97779 / GPL570 / 221562_s_at
GSE97779 / GPL570 / 221913_at
GSE97779 / GPL570 / 49327_at
control
arthritis
expression value
C
arthritis-control
adj.p < 0.05 & log2FC > 0
adj.p < 0.05 & log2FC < 0
adj.p >= 0.05
−log10(P.value)
Log2 fold change
SIRT3
EIF2AK2
IFIH1
OAS1
IFI44
ISG15
MX1
IFITM1
D
Gene: SIRT3
Gene: ISG15
Gene: MX1
Gene: IFI44
Expression
221562_s_at
221913_at
49327_at
205483_s_at
202086_at
214059_at
214453_s_at
control
arthritis

**Figure S4.**

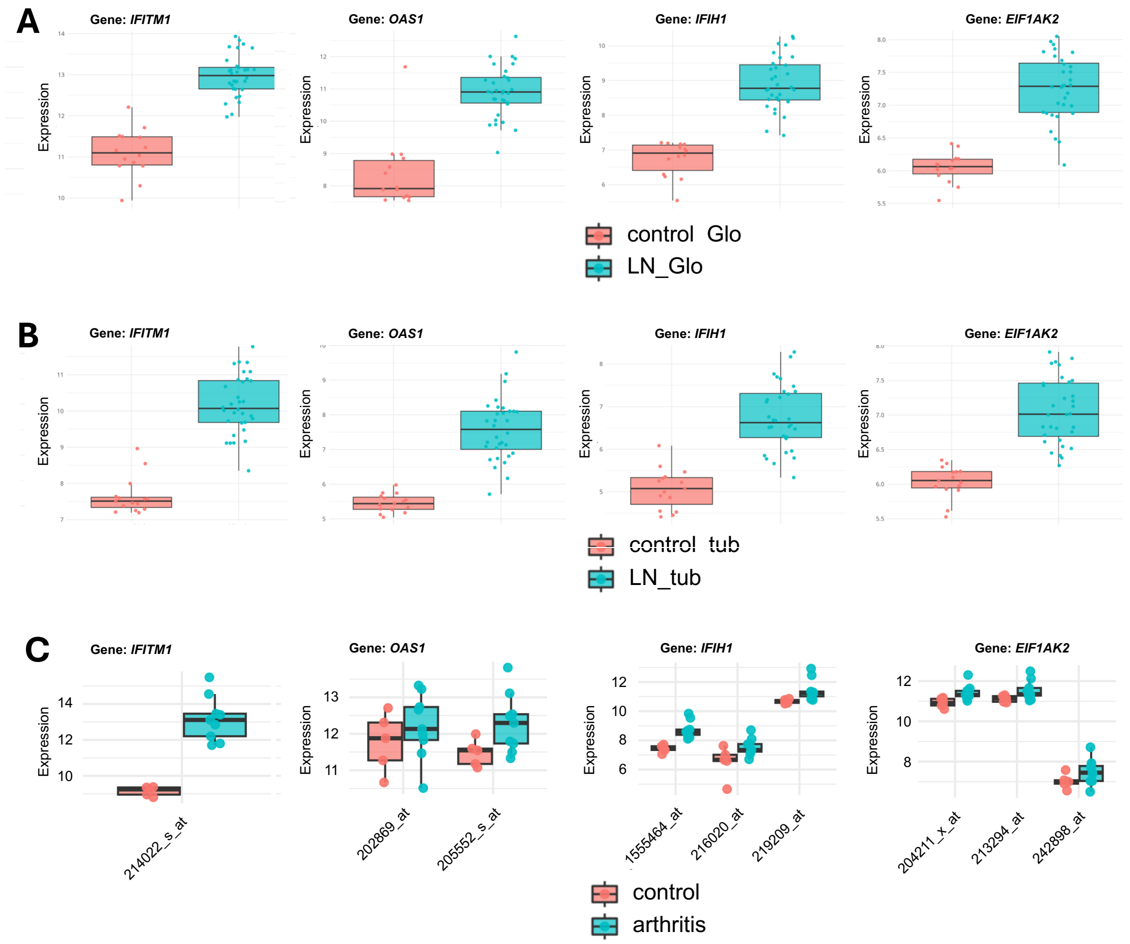
A
Gene: IFITM1
Gene: OAS1
Gene: IFIH1
Gene: EIF1AK2
Expression
control_Glo
LN_Glo
B
Gene: IFITM1
Gene: OAS1
Gene: IFIH1
Gene: EIF1AK2
Expression
control_tub
LN_tub
C
Gene: IFITM1
Gene: OAS1
Gene: IFIH1
Gene: EIF1AK2
Expression
214022_s_at
202869_at
205552_s_at
1555464_at
216020_at
219209_at
204211_x_at
213294_at
242898_at
control
arthritis
